# CHaRTr: An R toolbox for modeling Choices and Response Times in decision-making tasks

**DOI:** 10.1101/570184

**Authors:** Chandramouli Chandrasekaran, Guy E. Hawkins

## Abstract

Decision-making is the process of choosing and performing actions in response to sensory cues so as to achieve behavioral goals. A sophisticated research effort has led to the development of many mathematical models to describe the response time (RT) distributions and choice behavior of observers performing decision-making tasks. However, relatively few researchers use these models because it demands expertise in various numerical, statistical, and software techniques. Although some of these problems have been surmounted in existing software packages, the packages have often focused on the classical decision-making model, the diffusion decision model. Recent theoretical advances in decision-making that posit roles for “urgency”, time-varying decision thresholds, noise in various aspects of the decision-formation process or low pass filtering of sensory evidence, have proven to be challenging to incorporate in a coherent software framework that permits quantitative evaluations among these competing classes of decision-making models. Here, we present a toolbox — *Choices and Response Times in R*, or *CHaRTr* — that provides the user the ability to implement and test a wide variety of decision-making models ranging from classic through to modern versions of the diffusion decision model, to models with urgency signals, or collapsing boundaries. Earlier versions of *CHaRTr* have been instrumental in a number of recent studies of humans and monkeys performing perceptual decision-making tasks. We also provide guidance on how to extend the toolbox to incorporate future developments in decision-making models.

## 1. Introduction

Perceptual decision-making is the process of choosing and performing appropriate actions in response to sensory cues to achieve behavioral goals (Freedman and Assad, 2011; Gold and Shadlen, 2007; Hoshi, 2013; O’Connell et al., 2018; Shadlen and Kiani, 2013; Shadlen and Newsome, 2001). A sophisticated research effort in multiple fields has led to the formulation of cognitive process models to describe decision-making behavior (Donkin and Brown, 2018; Ratcliff et al., 2016). The majority of these models are grounded in the “sequential sampling” framework, which posits that decision-making involves the gradual accumulation of noisy sensory evidence over time until a bound (or criterion/threshold) is reached (Brunton et al., 2013; Forstmann et al., 2016; Gold and Shadlen, 2007; Hanks et al., 2014; Ratcliff and McKoon, 2008; Ratcliff et al., 2016; Shadlen and Kiani, 2013). Models derived from the sequential sampling framework are typically elaborated with various systematic and random components so as to implement assumptions and hypotheses about the underlying cognitive processes involved in perceptual decision-making (Diederich, 1997a; Ratcliff et al., 2016).

The most prominent sequential sampling model of decision-making is the diffusion decision model (DDM), which has an impressive history of success in explaining the behavior of human and animal observers (e.g., Ding and Gold, 2012a,b; Forstmann et al., 2016; Palmer et al., 2005; Ratcliff et al., 2016; Tsunada et al., 2016). However, recent studies propose alternative sequential sampling models that do not involve the integration of sensory evidence over time. Instead, novel sensory evidence is multiplied by an urgency signal that increases with elapsed decision time, and a decision is made when the signal exceeds the criterion (Cisek et al., 2009; Ditterich, 2006b; Thura et al., 2012). Another line of research proposes that observers aim to maximize their reward rate and suggests that the decision boundary dynamically decreases as the time spent making a decision grows longer. Such a framework has been argued to provide a better explanation for decision-making behavior in the face of sensory uncertainty (Drugowitsch et al., 2012).

One approach to distinguish between these different models is to systematically manipulate the stimulus statistics and/or the task structure and then test whether behavior is qualitatively consistent with one or another sequential sampling model (Brunton et al., 2013; Carland et al., 2015; Cisek et al., 2009; Scott et al., 2015; Thura and Cisek, 2014). An alternative approach is to quantitatively analyze the choice and RT behavior with a large set of candidate models, and then carefully use model selection techniques to understand the candidate models that best describe the data (e.g., Chandrasekaran et al., 2017; Evans et al., 2017; Hawkins et al., 2015b; Purcell and Kiani, 2016). The quantitative modeling and model selection approach allows the researcher to determine whether a particular model component (e.g., an urgency signal, or variability in the rate of information accumulation) is important for generating the observed behavioral data. It also provides a holistic method for testing model adequacy, because the proposed model is judged on its ability to account for all available data rather than focusing on a specific subset of the data (e.g., Evans et al., 2017).

Despite the apparent benefits of model selection, there are technical and computational challenges in the application of decision-making models to behavioral data. Some researchers have surmounted these issues by simplifying the process: using analytical solutions for the predicted mean RT and accuracy from the simplest form of the DDM, applied to participant-averaged behavioral data (Palmer et al., 2005; Tsunada et al., 2016). However, the complete distribution of RTs is highly informative, and often necessary, to reliably discriminate between the latent cognitive processes that influence decision-making (Forstmann et al., 2016; Luce, 1986; Ratcliff and McKoon, 2008; Ratcliff et al., 2016). Until recently, applying sequential sampling models like the DDM to the joint distribution over choices and RT required bespoke domain knowledge and computational expertise. This has hindered the widespread adoption of quantitative model selection methods to study decision-making behavior.

Some recent attempts have demystified the application of cognitive models of decision-making to behavioral data, providing a path for researchers to apply these methods to their own research questions. For instance, Vandekerckhove and Tuerlinckx developed the Diffusion Modeling and Analysis Toolbox (Vandekerckhove and Tuerlinckx, 2008), and Voss and Voss developed the diffusion model toolbox (fast-dm; Voss and Voss, 2007, 2008). Modern releases have improved the parameter estimation algorithms and can leverage multiple observers to perform hierarchical Bayesian inference (Wiecki et al., 2013). In hBayesDM (Ahn et al., 2017) and Dynamic Models of Choice (Heathcote et al., 2018) researchers can apply a range of models to behavior from a wide variety of decision-making paradigms ranging from choice tasks to reversal learning and inhibition tasks.

A common feature across all of the excellent toolboxes currently available is that they only provide optimization code to apply the DDM to data, or the DDM in addition to a few alternative models. As a consequence, the toolboxes provide no pathway for a researcher to rigorously compare the quantitative account of the DDM to alternative theories of the decision making process, including models with an urgency signal (Ditterich, 2006a), urgency-gating (Cisek et al., 2009), or collapsing bounds (Hawkins et al., 2015b). Simply put, we currently have no openly available and extensible toolbox for understanding choice and RT behavior using the many hypothesized models of decision-making. We believe there is a critical need for examining how these different models perform in terms of describing decision-making behavior.

The objective of this study was to address this need and provide a straightforward framework to analyze a range of existing sequential sampling models of decision-making behavior. Specifically, we aimed to provide an open-source and extensible framework that permits quantitative implementation and testing of novel candidate models of decision-making. The outcome of this study is *CHaRTr*, a novel toolbox in the R programming environment that can be used to analyze choice and RT data of humans and animals performing two-alternative forced choice tasks that involve perceptual or other types of decision-making. R is an open source language that enjoys widespread use and is maintained by a large community of researchers. *CHaRTr* can be used to analyze the RT and choice behavior of these observers, from the perspective of a (potentially large) range of decision-making models and can be readily extended when new models are developed. These new models can be incorporated into the toolbox with minimal effort and require only basic working knowledge of R and C programming; we explain the required skills in this manuscript. Similarly, new optimization routines that are readily available as R packages can be implemented if desired.

## 2. Methods and Materials

The methods are focused on the specification of various mathematical models of decision-making, and the parameter estimation and model selection processes. For reference, the symbols we use to describe the models are shown in Table 1. The naming convention for the models we have developed in *CHaRTr* is to take the main architectural feature of the model and use it as a prefix to the model. The diffusion decision model, henceforth DDM, refers to the simplest sequential sampling model, cDDM refers to a DDM with collapsing boundaries (Hawkins et al., 2015b), dDDM refers to a DDM with urgency signal defined by Ditterich (2006a), uDDM refers to a DDM with a linear urgency signal, and bUGM refers to an urgency gating model (UGM) with a linear urgency signal composed of a slope and an intercept (Thura et al., 2012).

**Table 1:** List of symbols used in the decision-making models implemented in *CHaRTr*

| Parameter | Description |
| --- | --- |
| $x(t)$ | State of the decision variable at time $t$. |
| $dt$ | Time step of the decision variable. |
| $z, s_z$ | Starting state of the decision variable (i.e., $x(0) = z$), and decision-to-decision variability in starting state. $s_z$ is the range of a uniform distribution with mean (midpoint) $z$. |
| $v_i, s_v$ | Rate at which the decision variable accumulates decision-relevant information (drift rate, $v$) in condition $i$, and decision-to-decision variability in drift rate. $s_v$ is the standard deviation of a normal distribution with mean $v_i$. |
| $\gamma(t)$ | Urgency signal that dynamically modulates the decision variable as a function of $t$. Can take different functional forms in different models. |
| $a_{upper}, a_{lower}$ | Upper and lower response boundaries that terminate the decision process. |
| $a_{upper}(t), a_{lower}(t)$ | Upper and lower response boundaries that vary as a function of $t$. |
| $T_{er}, s_t$ | Time required for stimulus encoding and motor preparation/execution (non-decision time), and decision-to-decision variability in non-decision time. $s_t$ is the range of a uniform distribution with mean (midpoint) $T_{er}$. |
| $s$ | Within-decision variability in the diffusion process. Represents the standard deviation of a normal distribution. By convention, set to a fixed value to satisfy a scaling property of the model. |
| $E(t)$ | Momentary sensory evidence at time $t$. |
| $\mathcal{N}(0, 1)$ | Normal distribution with zero mean and unit variance. |
| $\mathcal{U}(l_1, l_2)$ | Uniform distribution over the interval $l_1$ and $l_2$. |

### 2.1. Mathematical Models of Decision-Making

Sequential sampling models of decision-making assume that RT comprises two components (Ratcliff and McKoon, 2008; Ratcliff et al., 2016). The first component is the decision time, which encompasses processes such as the accumulation of sensory evidence and additional decision-related factors such as urgency. The second component is the non-decision time (or residual time), which involves the time required for processes that must occur to produce a response but fall outside of the decision-formation process, such as stimulus encoding, motor preparation and motor execution time.

We introduce various models of the decision-making process in approximately increasing level of complexity, beginning with the simple DDM.

#### 2.1.1. Simple Diffusion Decision Model (DDM)

The diffusion decision model (or DDM) is derived from one of the oldest interpretations of a statistical test – the sequential probability ratio test (Wald and Wolfowitz, 1948) – as a model of a cognitive process – how decisions are formed over time (Stone, 1960). The DDM provides the foundation for the decision-making models implemented in *CHaRTr* and assumes that decision-formation is described by a one-dimensional diffusion process (Fig. 1A) with the stochastic differential equation

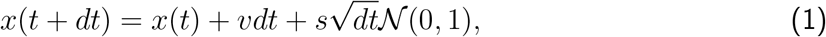

where *x*(*t*) is the state of the decision-formation process, known as the decision variable, at time *t*; *v* is the rate of accumulation of sensory evidence, known as the drift rate; *dt* is the step size of the process; *s* is the standard deviation of the moment-to-moment (Brownian) noise of the decision-formation process; 𝒩(0, 1) refers to a random sample from the standard normal distribution. A response is made when *x*(*t* + *dt*) ≥ *a_upper_* or *x*(*t* + *dt*) ≤ *a_lower_*. Whether a response is correct or incorrect is determined from the boundary that was crossed and the valence of the drift rate (i.e., *v >* 0 implies the upper boundary corresponds to the correct response, *v <* 0 implies the lower boundary corresponds to the correct response). In Fig. 1a, and in all DDM models in *CHaRTr*, we specify *a_lower_* = 0 and *a*_*upper*_ = *A*, without loss of generality. *z* represents the starting state of the evidence accumulation process (i.e., the position of the decision variable at *x*(0)) and can be freely estimated between *a*_*lower*_ and *a*_*upper*_. When we assume there is no a priori response bias, *z* is fixed to the midpoint between *a*_*lower*_ and *a*_*upper*_ (i.e., *A/*2). The decision time is the first time step *t* at which the decision variable crosses one of the two decision boundaries. The predicted RT is given as a sum of the decision time and the non-decision time *T*_*er*_.

**Figure 1:**
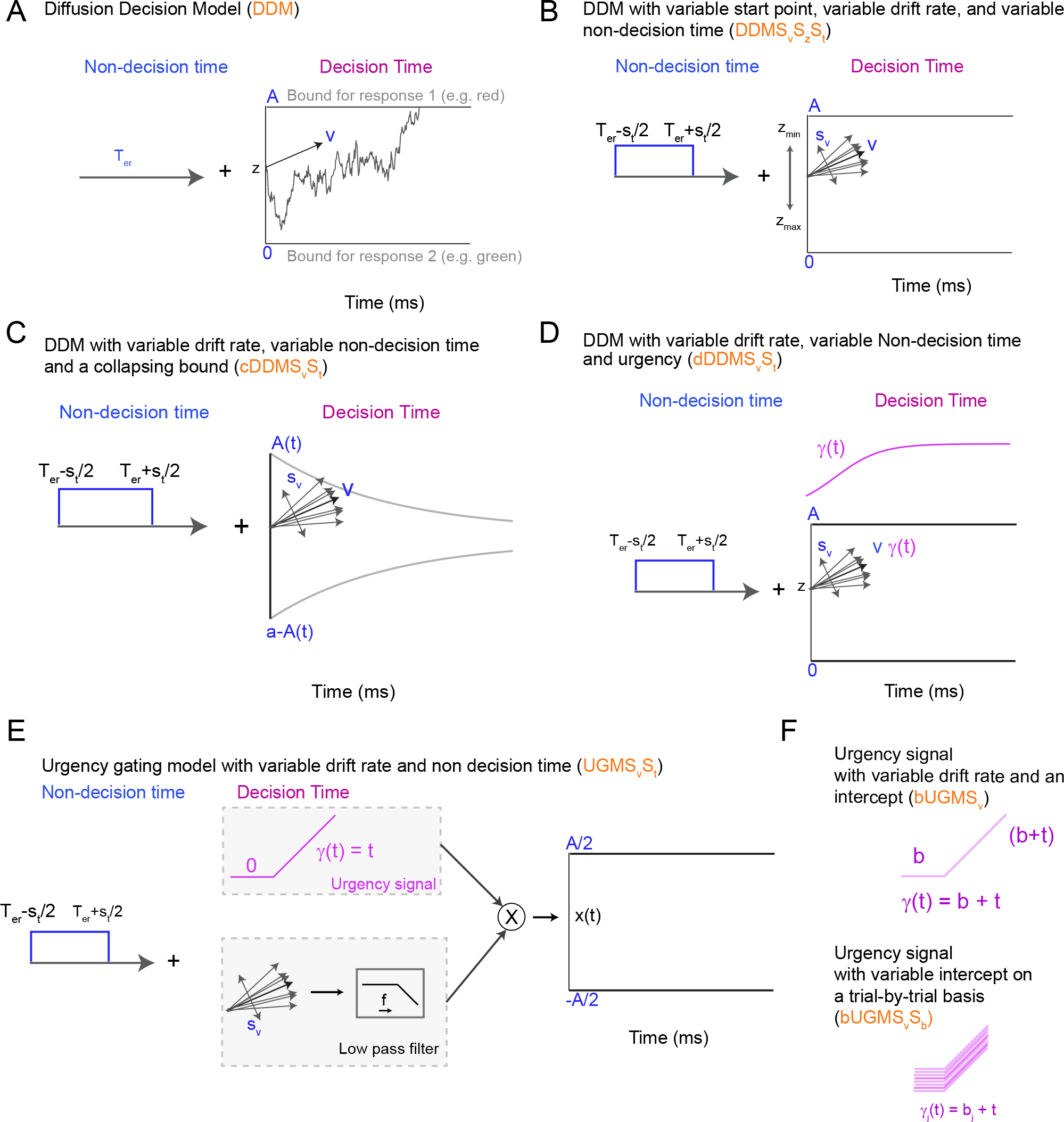
Schematic of some sequential sampling models of decision-making incorporated in *CHaRTr*. (A) The DDM model is the simplest example of a diffusion model of decision-making. (B) A variant of the DDM with variable non-decision time (*S*_*t*_), variable drift-rate (*S*_*v*_) and a variable start point (*S*_*z*_). (C) A DDM with collapsing bounds and variability in the non-decision time and drift rate. The function A(t) takes the form of a Weibull function as defined in Equation 6. (D) A variant of the DDM with variable non-decision time and drift rate, and an “urgency signal”. This urgency signal grows with elapsed decision time, which is implemented by multiplying the decision variable by the increasing function of time *γ*(*t*) (Equation 10, following Ditterich, 2006a). (E) UGM with variable drift rate (S_v_) and variable non decision time (S_t_). In the standard UGM, the urgency signal is only thought to depend on time and thus starts at 0. The sensory evidence is passed through a low pass filter (typically a 100-250 ms time constant, Carland et al., 2015; Thura et al., 2012). The sensory evidence is then multiplied by the urgency signal to lead to a decision variable that is then compared to boundaries. (F) Schematic of urgency signals with an intercept (top panel) and a variable intercept (bottom panel)

#### 2.1.2. DDM with Variable Starting State, Variable Drift Rate, and Variable Non-Decision Time

The (simple) DDM assumes a level of constancy from one decision to the next in various components of the decision-formation process: it always commences with the same level of response bias (*z*), the drift rate takes a single value (*v*_*i*_, for trials in experimental condition *i*), and the non-decision time never varies (*T*_*er*_).

None of these simplifying assumptions are likely to hold in experimental contexts. For example, the relative speed of correct and erroneous responses can differ, and participants’ arousal may exhibit random fluctuations over time, possibly due to a level of irreducible neural noise. Decades of research into decision-making models suggests that these effects, and others, are often well explained by combining systematic and random components in each of the starting state, drift rate, and non-decision time (Fig. 1B). In *CHaRTr*, we provide variants of the DDM where all of these parameters can be randomly drawn from their typically assumed distributions over different trials,

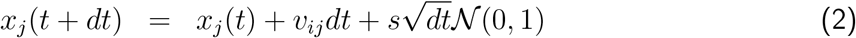

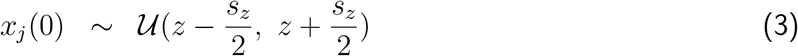

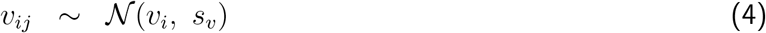

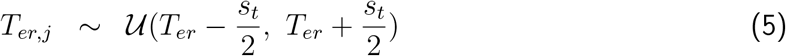

where *i* denotes an experimental condition; *j* denotes an exemplar trial; 𝒰 denotes the uniform distribution. *CHaRTr* provides flexibility to the user such that they can assume the decision-formation process involves none, some or all of these random components. Furthermore, it provides flexibility to assume distributions for the random components beyond those that have been typically assumed and studied in the literature. For example, one could hypothesize that non-decision times are exponentially distributed rather than uniformly distributed (Ratcliff, 2013).

#### 2.1.3 DDM with Collapsing Decision Boundaries (cDDM)

The DDM with collapsing boundaries generalizes the classic DDM by assuming that the sensory evidence required to commit to a decision is not constant as a function of elapsed decision time. Instead, it assumes that the decision boundaries gradually decrease as the decision-formation process grows longer and longer (e.g., Bowman et al., 2012; Drugowitsch et al., 2012; Hawkins et al., 2015a; Milosavljevic et al., 2010; Tajima et al., 2016). Collapsing boundaries terminate slower decisions based on weaker sensory signals (i.e., lower drift rates) at earlier time points than models with ‘fixed’ boundaries (i.e., simple DDM) and otherwise equivalent parameter settings. The net result of collapsing boundaries is a reduction in the positive skew (right tail) of the predicted RT distribution relative to the fixed boundaries DDM. This signature in the predicted RT distribution holds whether there is variability in parameters across trials (Section 2.1.2) or not (Section 2.1.1).

Collapsing boundaries allow the observer to implement a decision strategy where they do not commit an inordinate amount of time to decisions that are unlikely to be correct (i.e., decision processes with weak sensory signals). This allows the observer to sacrifice accuracy for a shorter decision time, so they can engage in new decisions that might contain stronger sensory signals and hence a higher chance of a correct response. When a sequence of decisions varies in signal-to-noise ratio from one trial to the next, like a typical difficulty manipulation in decision-making studies, collapsing boundaries are provably more optimal than fixed boundaries in the sense that they lead to greater predicted reward across the entirety of the decision sequence (Drugowitsch et al., 2012; Tajima et al., 2016). In this type of decision environment, collapsing boundaries have provided a better quantitative account of animal behavior, including monkeys, who might be motivated to obtain rewards to a greater extent than humans, possibly due to the operant conditioning and fluid/food restriction procedures used to motivate these animals (Hawkins et al., 2015a). Whether humans also aim to maximize reward is less clear.

Fig. 1C shows a schematic of a collapsing boundaries model. In *CHaRTr* we assume the collapsing boundary follows the cumulative distribution function of the Weibull distribution, following Hawkins et al. (2015a). The Weibull function is quite flexible and can approximate many different functions that one might wish to investigate, including the exponential and hyperbolic functions. We assume the lower and upper boundaries follow the form

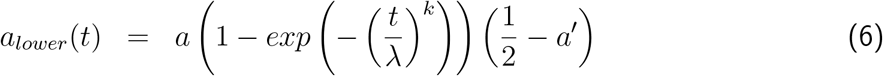

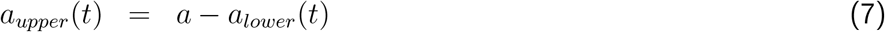

where *a*_*lower*_(*t*) and *a*_*upper*_(*t*) denote the position of the lower and upper boundaries at time *t*; *a* denotes the position of upper boundary at *t* = 0 (initial boundary setting, prior to any collapse); *a′* denotes the asymptotic boundary setting, or the extent to which the boundaries collapsed (the maximal possible collapse – where the upper and lower boundaries meet – can occur when *a′* = 1*/*2); *λ* and *k* denote the scale and shape parameters of the Weibull distribution.

The collapsing boundaries are denoted in *CHaRTr* as cDDM. When the *k* parameter is fixed to a particular value to aid stronger identifiability in parameter estimation (Hawkins et al., 2015a), we refer to the architecture as *cfk* to denote a fixed *k* value, here chosen to be 3 but can be modified in user implementations.

The collapsing boundaries, as implemented here, are symmetric, though they need not be; *CHaRTr* provides flexibility to modify all features of the boundaries, including symmetry for each response option, and the functional form. For instance, one might hypothesize that linear collapsing boundaries are a better description of the decision-formation process than nonlinear boundaries (Murphy et al., 2016; O’Connell et al., 2018). *CHaRTr* also permits DDMs with collapsing boundaries to incorporate any combination of variability in starting state, drift rate, and non-decision time (e.g., models of the form cDDMS_v_S_z_S_t_ and cfkDDMS_v_S_z_S_t_).

#### 2.1.4. DDM with an Urgency Signal (uDDM)

The DDM with an urgency signal assumes that the input evidence – consisting of the sensory signal and noise – is modulated by an “urgency signal”. This urgency-modulated sensory evidence is accumulated into the decision variable throughout the decision-formation process. As the process takes longer, the urgency signal grows in magnitude, implying that sensory evidence arriving later in the decision-formation process has a more profound impact on the decision-variable than information arriving earlier (Fig. 1D). To make the distinction between an urgency signal and collapsing boundaries clear, the DDM with an urgency signal assumes a dynamically modulated input signal combined with boundaries that mirror those in the classic DDM; the DDM with collapsing boundaries assumes a decision variable that mirrors the classic DDM combined with dynamically modulated decision boundaries.

As with the collapsing boundaries, the urgency signal can take many functional forms; we have implemented two such forms in *CHaRTr*. The general implementation of the urgency signal is

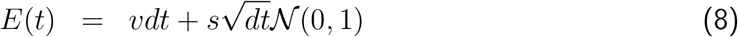

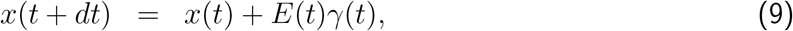

where *E*(*t*) denotes the momentary sensory evidence at time *t*; *γ*(*t*) denotes the magnitude of the urgency signal at time *t*. Note that with increasing decision time the urgency signal magnifies the effect of the sensory signal (*vdt*) and the sensory noise 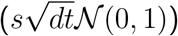.

The first urgency signal implemented in *CHaRTr* follows a 3 parameter logistic function with two scaling factors (*s*_*x*_, *s*_*y*_) and a delay (*d*), originally proposed by Ditterich (2006a):

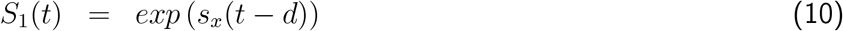

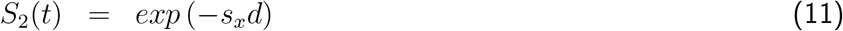

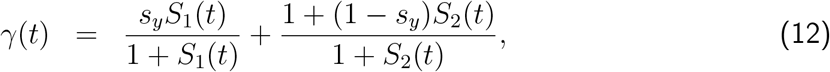

The second form of urgency signal implemented in *CHaRTr* follows a simple, linearly increasing function

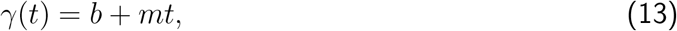

where *b* is the intercept of the urgency signal. The slope is assumed to be m. As with the DDMs described above, urgency signal models can incorporate any combination of variability in starting state, drift rate and non-decision time, giving rise to a family of different decision-making models. We also allow for the possibility of variability across decisions in the intercept term of the linear urgency signal,

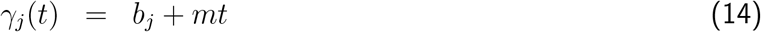

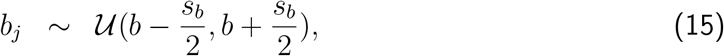

where *j* denotes an exemplar trial, and *b* and *s*_*b*_ denote the mean (i.e., midpoint) and range of the uniform distribution assumed for the urgency signal.

In *CHaRTr*, we have assumed that the urgency signal exerts a multiplicative effect on the sensory evidence (Equation 9). One variation of urgency signal models proposed in the literature posits that urgency is added to the sensory evidence, rather than multiplied by it (Hanks et al., 2014, 2011). In the one-dimensional diffusion models considered here, additive urgency signals make predictions that cannot be discriminated from a DDM with collapsing boundaries (Boehm et al., 2016). That is, for any functional form of an additive urgency signal, there is a function for the collapsing boundaries that will generate identical predictions. For this reason we do not provide an avenue for simulating and estimating additive urgency signal models in *CHaRTr*, and instead recommend the use of the DDM with collapsing boundaries.

#### 2.1.5 Urgency Gating Model (UGM)

In a departure from the classic DDM framework, the Urgency Gating Model (UGM) proposes there is no integration of evidence, at least not in the same form as the DDM (Cisek et al., 2009; Thura et al., 2012; Thura and Cisek, 2014). Rather, the UGM assumes that incoming sensory evidence is low-pass filtered, which prioritizes recent over temporally distant sensory evidence, and this low-pass filtered signal is modulated by an urgency signal that increases linearly with time (Equation 13).

Implementation of the UGM in *CHaRTr* uses the exponential average approach for discrete low-pass filters (smoothing). The momentary evidence for a decision is a weighted sum of past and present evidence, which gives rise to the UGM’s pair of governing equations

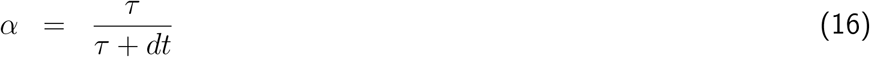

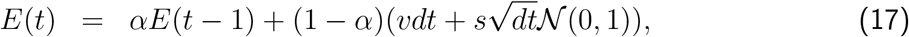

where *τ* is the time constant of the low-pass filter, which has typically been set to relatively small values of 100 or 200 ms in previous applications of the UGM, and *α* controls the amount of evidence from previous time points that influences the momentary evidence at time *t*. For instance, when *α* = 0 there is no low-pass filtering, and when *τ* = 100*ms* (and *dt* is 1 ms) the previous evidence is weighted by 0.99 and new evidence by 0.01.

The decision variable at time *t* is now given as

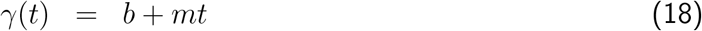

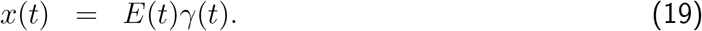

The intercept and slope of the urgency signal are set to particular values in standard applications of the UGM (*b* = 0, *m* = 1), reducing equation 19 to

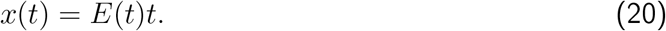

In *CHaRTr*, we allow for variants of the UGM where the parameters of the urgency signal are not fixed. For instance, similar to the DDM with an urgency signal, we can test a UGM where the intercept (*b*) is freely estimated from data (bUGM), and even an intercept that varies on a trial-by-trial basis (Equation 14).

### 2.2 Fitting Models to Data: Parameter Estimation and Model Selection

#### 2.2.1. Parameter Estimation

In *CHaRTr*, we estimate parameters for each model and participant independently, using Quantile Maximum Products Estimation (QMPE; Heathcote and Brown, 2004; Heathcote et al., 2002). QMPE uses the QMP statistic, which is similar to *χ*^2^ or multinomial maximum likelihood estimation and quantifies agreement between model predictions and data by comparing the observed and predicted proportions of data falling into each of a set of inter-quantile bins. These bins are calculated separately for the correct and error RT data. In all examples that follow, we use 9 quantiles calculated from the data (i.e., split the RT data into 10 bins), though the user can specify as many quantiles as they wish. Generally speaking, we recommend no fewer than 5 quantiles, to prevent loss of distributional information, and no more than approximately 10 quantiles, to prevent noisy observations in observed data especially at the tails of the distribution potentially bearing undue influence on the parameter estimation routine.

Many of the models considered in *CHaRTr* have no closed-form analytic solution for their predicted distribution. To evaluate the predictions of each model, we simulate 10,000 Monte Carlo replicates per experimental condition during parameter estimation. Once the parameter search has terminated, we use 50,000 replicates per experimental condition to precisely evaluate the model predictions and perform model selection. In *CHaRTr*, the user can vary the number of replicates used for parameter estimation and model selection; in previous applications, we have found these default values provide an appropriate balance between precision of the model predictions and computational efficiency. To simulate the models, we use Euler’s method, which approximates the models’ representation as stochastic differential equations.

Alternatives to our simulation-based approach exist, such as the integral equation methods of Smith (2000) or others that use analytical techniques to calculate first passage times (Gondan et al., 2014; Navarro and Fuss, 2009), to generate exact distributions. We do not pursue those methods in *CHaRTr* owing to the model-specific implementation required, which is inconsistent with *CHaRTr*’s core philosophy of allowing the user to rapidly implement a variety of model architectures.

We estimate the model parameters using differential evolution to optimize the goodness of fit (DEoptim package in R, Mullen et al., 2011). For the type of non-linear models considered in *CHaRTr*, we have previously found that differential evolution more reliably recovers the true data generating model than particle swarm and simplex optimization algorithms (Hawkins et al., 2015a). DEoptim also allows easy parallelization and can be used readily in the cloud with large number of cores to speed the process of model estimation. However, we again provide flexibility in this respect; the user can change this default setting and specify their preferred optimization algorithm/s.

#### 2.2.2 Model Selection

*CHaRTr* provides two metrics for quantitative comparison between models. Each metric is based on the maximized value of the QMP statistic, which is a goodness-of-fit term that approximates the continuous maximum likelihood of the data given the model.

The DDM is a special case of the model variants considered and will almost always fit more poorly than any of the other variants. We provide model selection methods that determine if the incorporation of additional components such as urgency or collapsing bounds provide an improvement in fit that justifies the increase in model complexity.

The raw QMP statistic, as an approximation to the likelihood, can be used to calculate the Akaike Information Criterion (Akaike, 1974, AIC) and the Bayesian Information Criterion (Schwarz, 1978, BIC). We provide methods to compute AIC and BIC owing to the differing assumptions underlying the two information criteria (Aho et al., 2014), and differing performance with respect to the modeling goal (Evans, in press).

*CHaRTr* also provides functionality to transform the model selection metrics into model weights, which account for uncertainty in the model selection procedure and aid interpretation by transformation to the probability scale. The weight *w* for model *i*, *w*(*M*_*i*_), relative to a set of *m* models, is given by

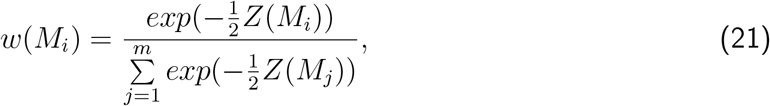

where *Z* is AIC, BIC, or the deviance (−2× log-likelihood; that is, −2× QMP statistic). The model weight is interpreted differently depending on the metric *Z*:

- Where *Z* is the log-likelihood, the model weights are relative likelihoods. *Z* should only be used in the model weight transformation when all models under consideration have the same number of freely estimated parameters.
- Where *Z* is the AIC, the model weights become Akaike weights (Wagenmakers and Farrell, 2004).
- Where *Z* is the BIC, and the prior probability over the *m* models under consideration is uniform (i.e., each model is assumed to be equally likely before observing the data), the model weights approximate posterior model probabilities (*p*(*M |Data*), Wasserman, 2000).

#### 2.2.3 Visualization: Quantile Probability Plots

Visualization of choice and RT data is critical to understanding observed and predicted behavior. Such visualization can prove challenging in studies of rapid decision-making because each cell of the experimental design (e.g., a particular stimulus difficulty) yields a joint distribution over the probability of a correct response (accuracy) and separate RT distributions for correct and error responses. Since most decision-making tasks manipulate at least one experimental factor across multiple levels, such as stimulus difficulty, each data set is comprised of a family of joint distributions over choice probabilities and pairs of RT distributions (correct, error). Following convention and recommendation (Ratcliff and McKoon, 2008; Ratcliff et al., 2016), we visualize these joint distributions with quantile probability (QP) plots. QP plots are a compact form to display choice probabilities and RT distributions across multiple conditions.

In a typical QP plot, quantiles of the RT distribution of a particular type (e.g., correct responses) are plotted as a function of the proportion of responses of that type. For example, consider a hypothetical decision-making experiment with three different levels of stimulus difficulty; Fig. 2 provides a plausible example of the data from such an experiment. Now assume that for one of the experimental conditions, the accuracy of the observer was 55%. To display the choice probabilities, correct RTs and error RTs for this condition, the QP plot shows a vertical column of *N* markers above the *x*-axis position ~ 0.55, where the *N* markers correspond to the *N* quantiles of the RT distribution of correct responses (rightmost gray bar in Fig. 2). The QP plot also shows a vertical columns of *N* markers at the position 1 − 0.55 = 0.45, where this set of *N* markers correspond to the *N* quantiles of the distribution of error RTs (leftmost gray bar in Fig. 2). This means that RT distributions shown to the right of .5 on the *x*-axis reflect correct responses, and those to the left of .5 on the *x*-axis reflect error responses.

**Figure 2:**
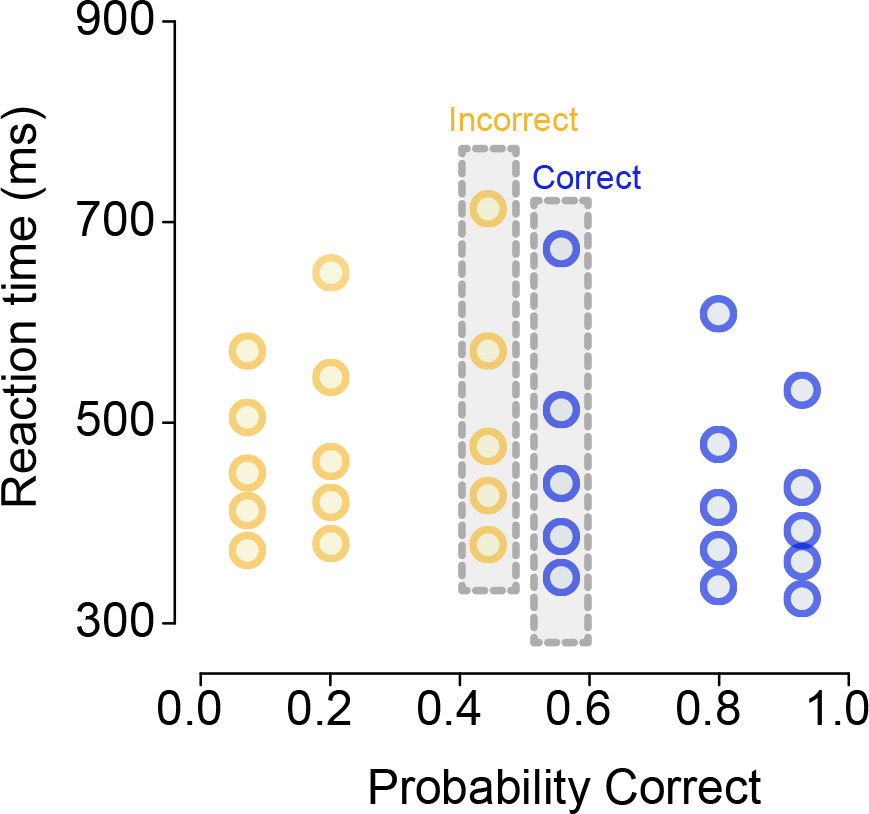
A quantile probability (QP) plot of choice and RT data from a hypothetical decision-making experiment with three levels of stimulus difficulty. The three difficulty levels are represented as vertical columns mirrored around the midpoint of the *x*-axis (.5). In this example, the lowest accuracy condition had ~ 55% correct responses, so the RTs for correct responses in this condition are located at .55 on the *x*-axis and the corresponding RTs for error responses are located at 1 −.55 = .45 on the *x*-axis; these two RT distributions are highlighted in gray bars. For each RT distribution we plot along the *y*-axis the 10^th^, 30^th^, 50^th^, 70^th^, 90^th^ percentiles (i.e., .1, .3, .5, .7, .9 quantiles), separately for correct and error responses in each of the three difficulty levels. For clarity, correct responses are shown in blue and error responses are shown in yellow.

The default *CHaRTr* QP plot display 5 quantiles of the RT distribution: 0.1, 0.3, 0.5, 0.7 and 0.9 (sometimes also referred to as five percentiles: 10^th^, 30^th^, 50^th^, 70^th^, 90^th^). The .1 quantile summarizes the leading edge of the RT distribution, the .5 quantile (median) summarizes the central tendency of the RT distribution, and the .9 quantile summarizes the tail of the RT distribution. The goal of visualization with QP plots, or other forms of visualization, is to enable comparison of the descriptive adequacy of a model’s predictions relative to the observed data.

## 3. Results

The results section first provides guidance on the use of *CHaRTr* and how to apply the various models of the decision-making process to data. The second part of the results section illustrates the use of *CHaRTr* to analyze RT and choice data from hypothetical observers, followed by a case study modeling data from two non-human primates (Roitman and Shadlen, 2002). Code for the *CHaRTr* toolbox is available at chartr.chandlab.org/ and will eventually be released as an R library.

### 3.1. Toolbox flow

Fig. 3 and Fig. 4 provide flowcharts for *CHaRTr*. Fig. 3 provides an overview of the five main steps involved in the cognitive modeling process. Fig. 4 provides a schematic overview of the steps involved in the parameter estimation component of the process, which uses the differential evolution optimization algorithm (Mullen et al., 2011).

**Figure 3:**
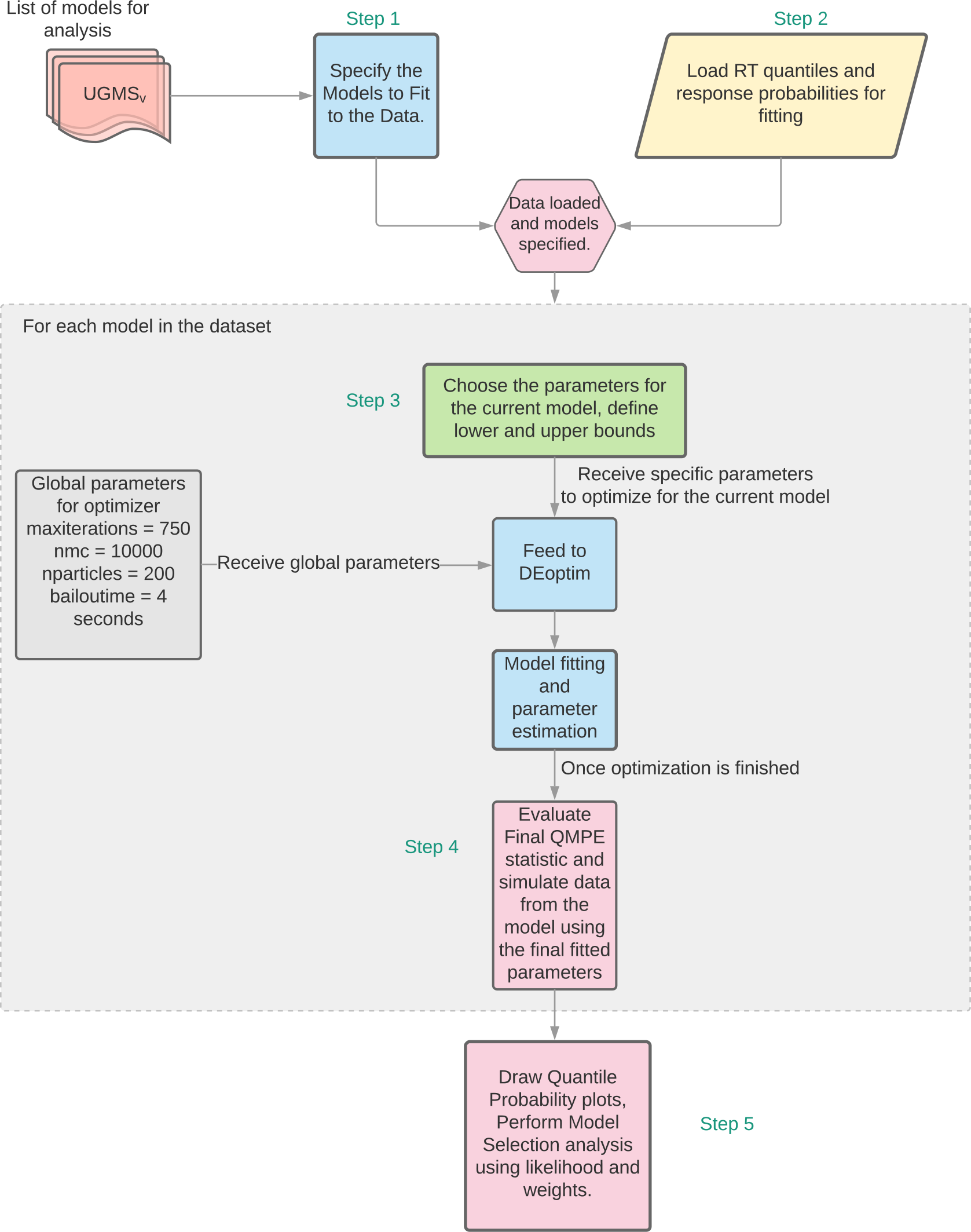
*CHaRTr* flow chart. Models are specified and once data is available, the parameters are estimated through the optimization procedure. Once parameter estimation is complete, the final goodness of fit statistic is calculated for every model under consideration, which is used for subsequent model selection analyses.

**Figure 4:**
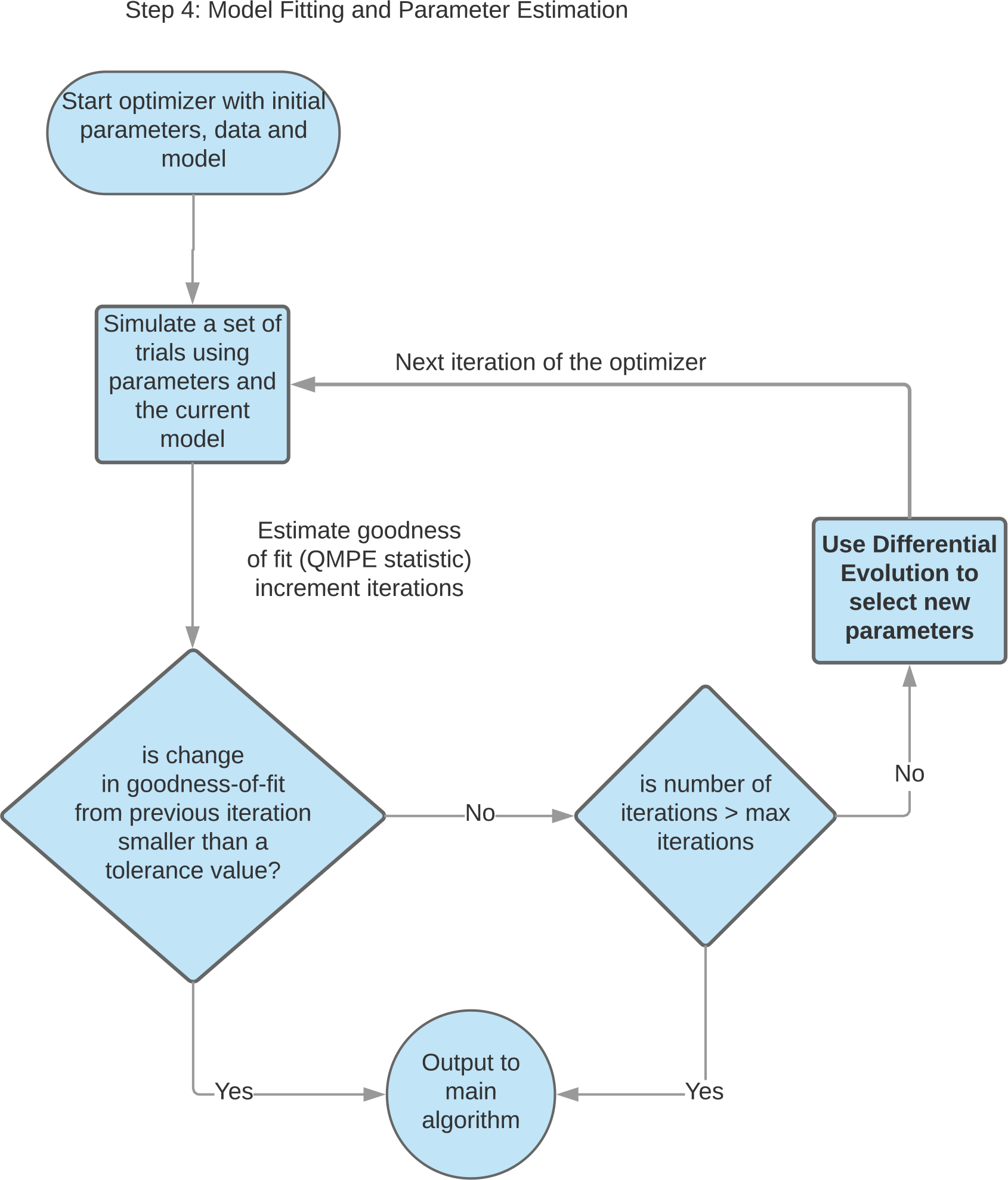
Flow chart for the parameter estimation component of *CHaRTr*, which uses the differential evolution optimization algorithm (Mullen et al., 2011).

The typical steps in *CHaRTr* for estimating the parameters of a decision-making model from data are as follows:

Step 1: **Model Specification**: Specify models in the C programming language, and compile the C code to create the shared object, chartr-modelspec.so, that is dynamically loaded into the R workspace. Future versions of *CHaRTr* will use the Rcpp framework and will not require the compilation and loading of shared objects (Eddelbuettel and François, 2011).

Step 2: **Formatting and Loading Data**: Convert raw data into an appropriate format (choice probabilities, quantiles of RT distributions for correct and error trials). Save this data object for each unit of analysis (e.g., a participant, different experimental conditions for the same participant). Load this data object into the R workspace.

Step 3: **Parameter Specification**: Choose the parameters of the desired model that need to be estimated along with lower and upper boundaries on those parameters (i.e., the minimum and maximum value that each parameter can feasibly take).

Step 4: **Parameter Estimation**: Pass the parameters, model and data to the optimization algorithm (differential evolution). The algorithm iteratively selects candidate parameter values and evaluates their goodness of fit to data. This process is repeated until the goodness of fit no longer improves (Fig. 4).

Step 5: **Model Selection**: The parameter estimates from the search termination point (i.e., the point where goodness of fit no longer improves), the corresponding goodness of fit statistics and model predictions are saved for subsequent model selection and visualization.

These 5 steps are repeated for each model and each participant under consideration. In the next few sections, we elaborate on each of the steps with examples to illustrate their implementation in *CHaRTr*. We note that use of *CHaRTr* requires a basic knowledge of R programming, and if one wishes to design and test a new decision-making model then also C programming. Owing to the many excellent online resources for both languages (a simple search of “R program tutorial” will return many helpful results), we do not provide a tutorial for either language here.

### 3.1.1 Model Specification

The difference equation for the model variants implemented in *CHaRTr* is specified in C code in the file “chartr-ModelSpec.c”. An example implementation of the DDM (Section 2.1.1) is shown in Listing 1. The functions take as input the various parameters that are to be optimized along with various constants such as the maximum number of time points to simulate as well as the time step.

#### Listing 1: Source code for a function in C that implements the difference equation for the DDM.

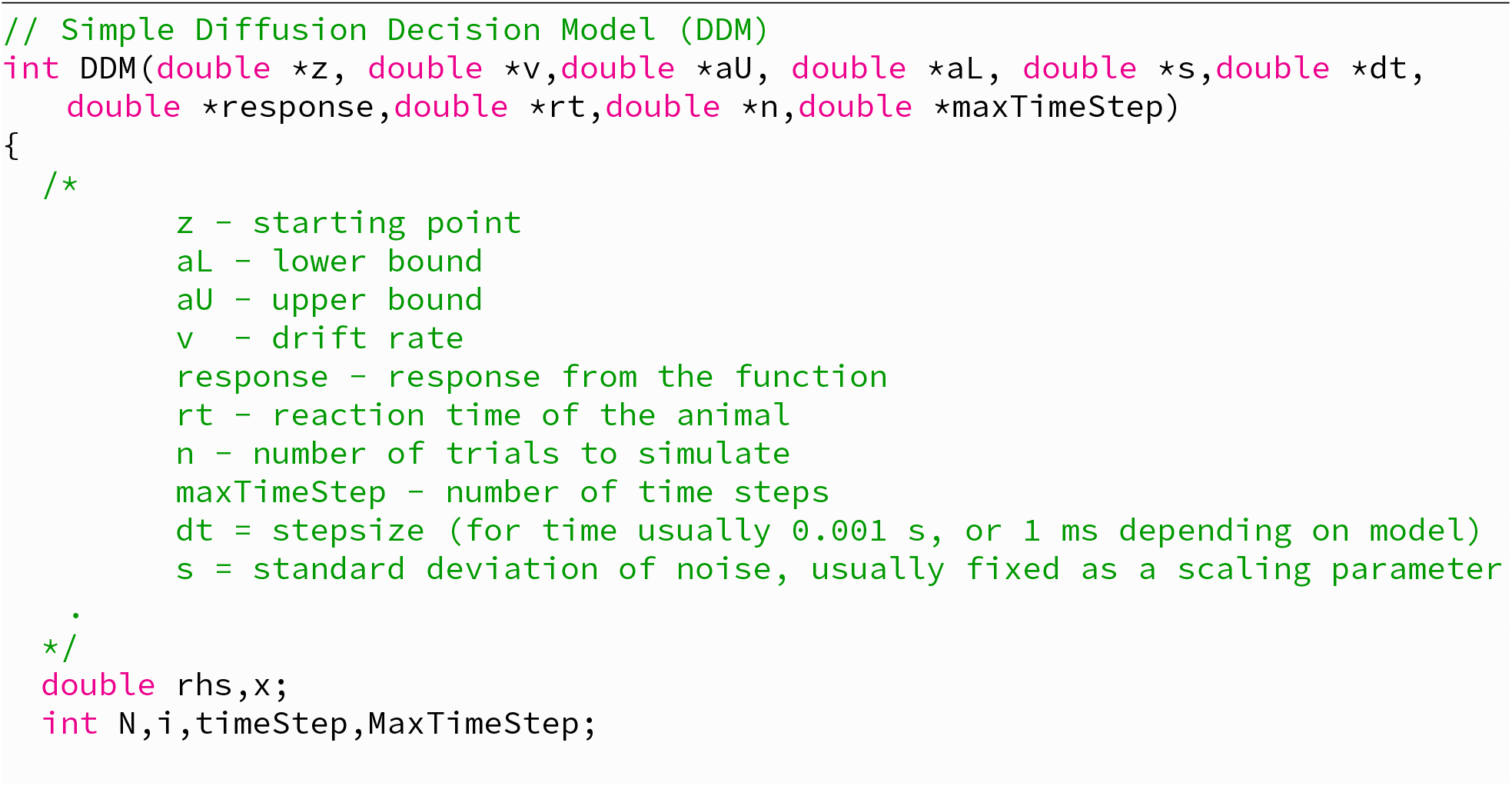
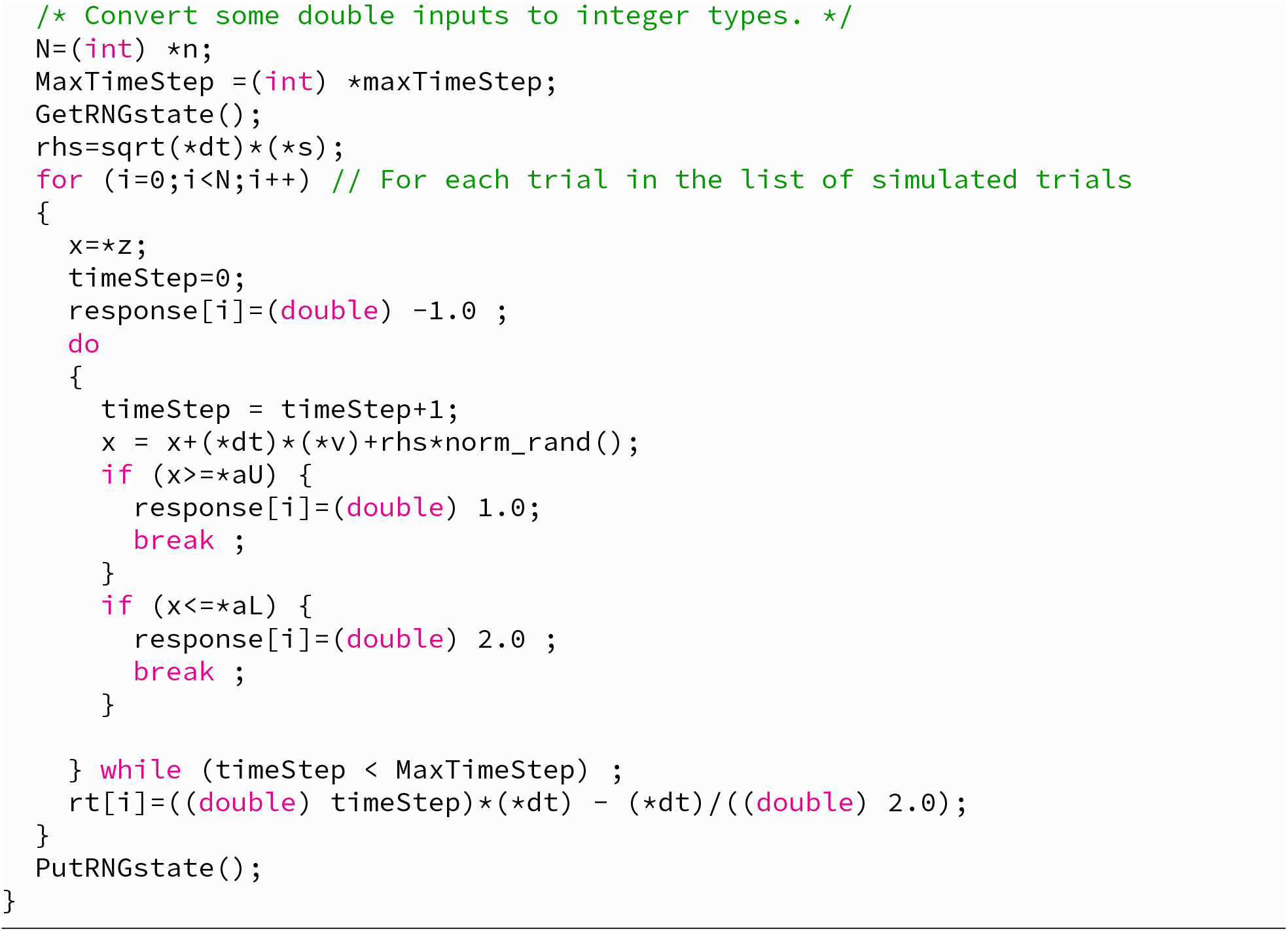

Once the C code has been specified for the model, the code is compiled using the following command that uses the SHLIB framework (R Core Team) at the terminal (usually ITERM in mac, Terminal Emulator in linux). The command shown in Listing 2 calls the appropriate compiler (clang on mac, gcc on linux), identifies the appropriate compiler to run, and loads the appropriate libraries and ensures the correct options are applied during compilation to create the architecture specific shared object.

#### Listing 2: Creating a shared library for loading the specified models into R.

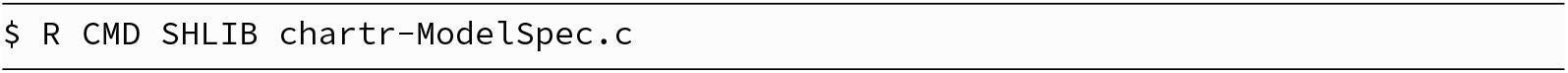

The typical output of the command when run successfully on Linux is shown in Listing 3. The output of the compilation is a shared object called *chartr-modelspec.so* that is dynamically loaded into R for use with the differential evolution optimizer.

#### Listing 3: Output from the compilation of the command in Listing 2.

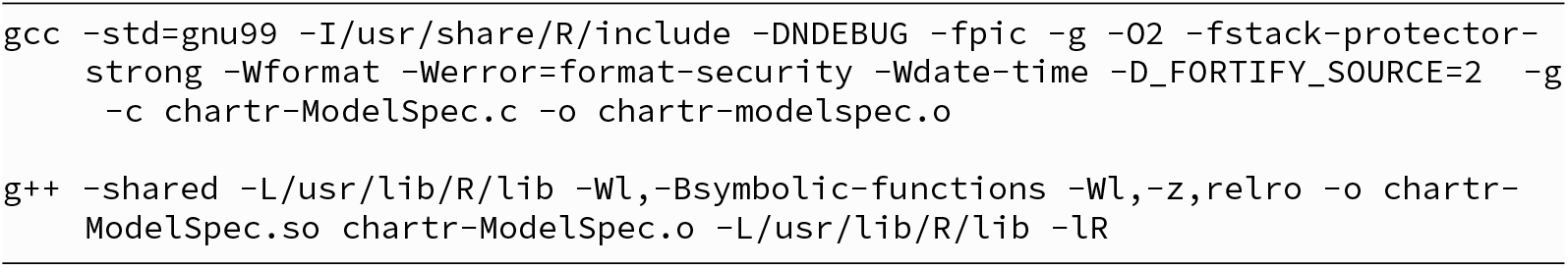

#### Listing 4: The required raw data format for parameter estimation in *CHaRTr*.

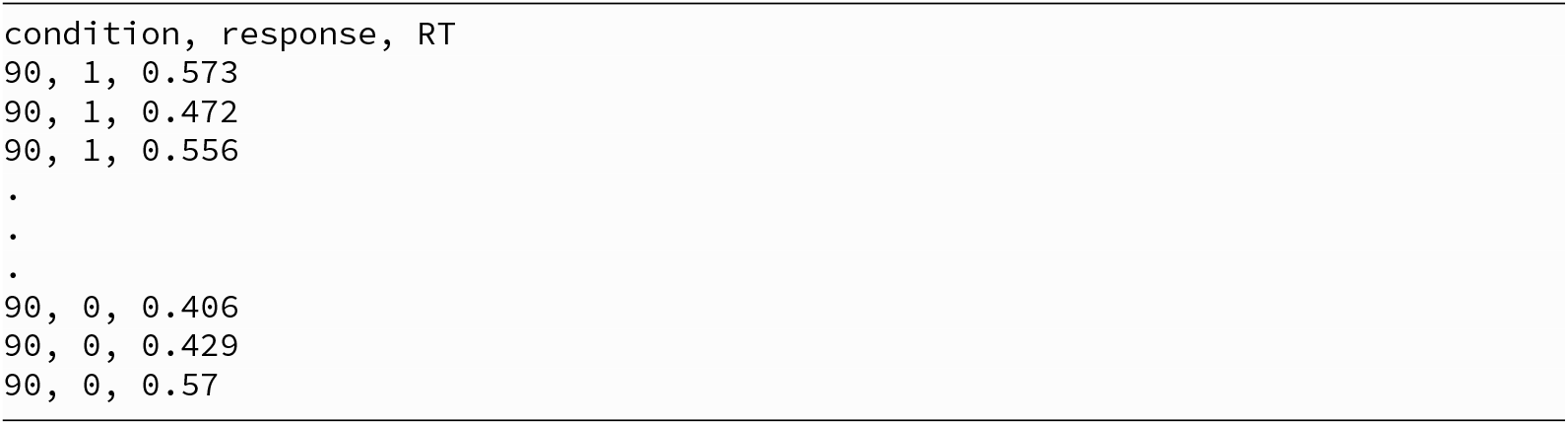

#### Listing 5: Loading data for “Subj1” for model estimation.

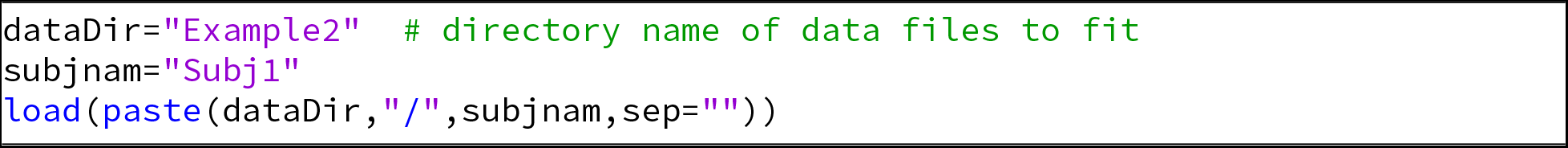

We anticipate that future versions of *CHaRTr* will use the Rcpp framework (Eddelbuettel and François, 2011), which will obviate the need for compiling and loading shared object libraries.

#### 3.1.2 Formatting and Loading Data

To estimate the parameters of decision-making models in *CHaRTr*, the data need to be organized in a separate comma separated values (CSV) file for each participant in a simple three column format: “condition, response, RT”. “condition” is typically a stimulus difficulty parameter, “response” is correct (1) or incorrect (0), and RT is the response time (or reaction time when response time and movement can be separated). For example, in a typical file, data for a single stimulus difficulty (e.g., one level of motion coherence in a random dot kinematogram) would look like Listing 4.

The raw data are converted in “chartr-processRawData.r” to generate 9 quantiles (10 bins) of correct and error RTs to be used in the parameter estimation process. It also stores the data as a R list named *dat*, which includes four fields: *n, p, q, pb*.

- *n* is the number of correct and error responses in each condition.
- *p* is the proportion of correct responses in each condition (derived from *n*).
- *q* is the quantiles of the correct and error RT distributions in each condition.
- *pb* is the number of responses in each bin of the correct and error RT distributions in each condition (derived from *n*).

*dat* is saved to disk as a new file. The *dat* file is loaded into the R workspace as required for the model estimation procedure. Listing 5 shows R code for loading RT and choice data, as stored in *dat*, for a given participant.

#### 3.1.3. Parameter Specification

The next step in model estimation is, for each model, to specify a list of parameters that can be freely estimated from data along with each parameter’s lower and upper bound; we provide default suggestions for the lower and upper boundaries in *CHaRTr*. Model parameters can be generated by calling the function *paramsandlims* with two arguments: model name and the number of stimulus difficulty levels in the experiment. The number of stimulus difficulties is internally converted into drift rate parameters; for example, if there are *n* stimulus difficulties, then *paramsandlims* will estimate *n* independent drift rate parameters. There is also functionality in *CHaRTr* to specify fixed (non-estimated) values of some parameters, such as a drift rate of 0 for conditions with non-informative sensory information (e.g., 0% coherence in a random dot kinematogram experiment). *paramsandlims* returns a named list with the following fields: lowers, uppers, parnames, fitUGM. These variables are used internally in the parameter estimation routines.

#### 3.1.4. Parameter Estimation

Steps 1–3 loaded the required data, identified the desired model to fit and specified the parameters of the model to be estimated. This information is now passed to the optimization algorithm (differential evolution). Parameter optimization is an iterative process of proposing candidate parameter values, accepting or rejecting candidate parameter values based on their goodness of fit, and repeating. This process continues until the proposed parameter values no longer improve the model’s goodness of fit. These are assumed to be the best-fitting parameter values, or the (approximate) maximum likelihood estimates. Fig. 4 provides an overview of the steps involved in parameter estimation when using the differential evolution optimization algorithm (Mullen et al., 2011).

The accompanying file “chartr-DemoFit.r” provides a complete code example for estimating the parameters of a model with urgency.

#### 3.1.5. Model Selection

Once the best-fitting parameters have been estimated from a set of candidate models, the final step is to use this information to guide inference about the relative plausibility of each of the models given the data. Many different levels of questions can be asked of these models. The best practices for model selection are described generally in Aho et al. (2014) and for the specific problem of behavioral modeling in Heathcote et al. (2015).

In *CHaRTr*, we provide functions for converting from the raw QMP statistic that approximates the likelihood. The likelihood provides essentially a goodness-of-fit statistic that can be combined with penalized model comparison metrics. This could entail comparison between two models at multiple levels of granularity. For instance, the question could be, “which of the models considered provides the better description of the data”, or “is a DDM with variable baseline better than a DDM without a variable baseline”. It could also be used to compare between a model with collapsing boundaries and a model with drift-rate variability (O’Connell et al., 2018) or between models with different forms of collapsing boundaries (Hawkins et al., 2015a). All of these questions can be answered using *CHaRTr*. As a guide, we provide illustrations of model selection analyses using *CHaRTr* in two case studies presented in Section 3.4. We also apply the model selection analyses to the behavior of monkeys performing a decision-making task (Roitman and Shadlen, 2002).

##### Listing 6: R Code for estimating the RTs and choice for the model DDM S_v_S_z_S_t_

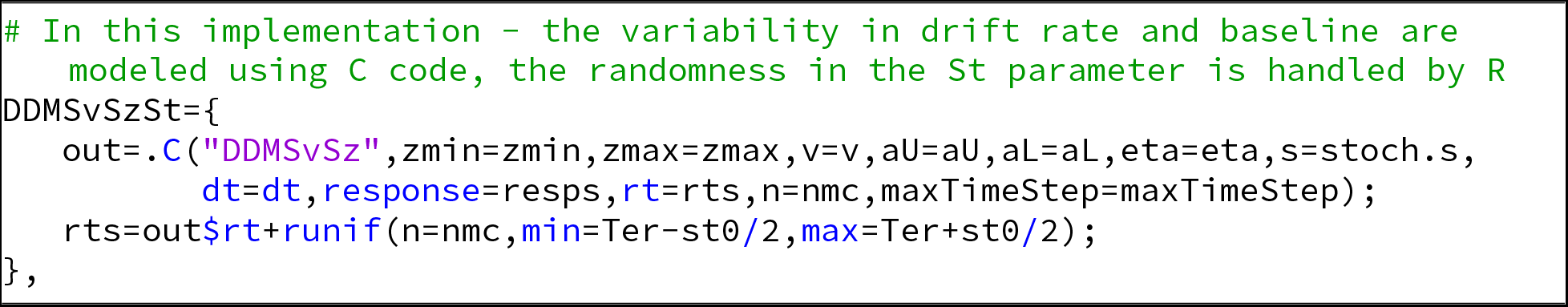

### 3.2 Extending CHaRTr

*CHaRTr* is designed with the goal of being readily extensible, to allow the user to specify new models with minimal development time. This allows the user to focus on the models of scientific interest while *CHaRTr* takes care of the model estimation and selection details behind the scenes. Here, we provide an overview of the steps required to add new models to *CHaRTr*.

1. Add a new function to “chartr-ModelSpec.c” with the parameters needed to be estimated for the model. Specify the model in C code, similar to Listing 1. Provide the new model with a unique name (i.e., not shared with any other models in the toolbox), preferably using the convention we defined above.
2. Add any new parameters of the model to the function *makeparamlist*, and to the *paramsandlims* function in script “chartr-HelperFunctions.r”.
3. Add the name of the model to the function *returnListOfModels*, in script “chartr-HelperFunctions.r”.
4. Make sure additional parameters are passed to the functions *diffusionC* and *getpreds*, in scripts “chartr-HelperFunctions.r” and “chartr-FitRoutines.r”, respectively.
5. Finally, specify in function *diffusionC* the code for generating RTs and responses to use for model fitting. For example, the code for generating the RTs and responses for DDMS_v_S_z_S_t_ is shown in the Listing 6

### 3.3. Simulating Data from Models in *CHaRTr*

Once models are specified, they can be used to generate simulated RTs and discrimination accuracy for each condition. Simulated data help refine quantitative hypotheses. They also provide much greater insight into the dynamics of different decision-making models and how different variables in these models modulate the predicted RT distributions for correct and error trials (Ratcliff and McKoon, 2008).

*CHaRTr* provides straightforward methods to simulate data from decision-making models and generate quantile probability plots to compactly summarize and visualize RT distributions and accuracy. The function *paramsandlims*, used above in the parameter estimation routine, can also be used to generate hypothetical parameters to be passed to the function *simulateRTs*, which generates a set of simulated RTs and choice responses. By hypothetical parameters, we mean a set of reasonable starting values. An example is shown in Listing 7. These parameters can be changed by the user.

#### Listing 7

R code for simulating RT and choice responses from the simple diffusion decision model (DDM).

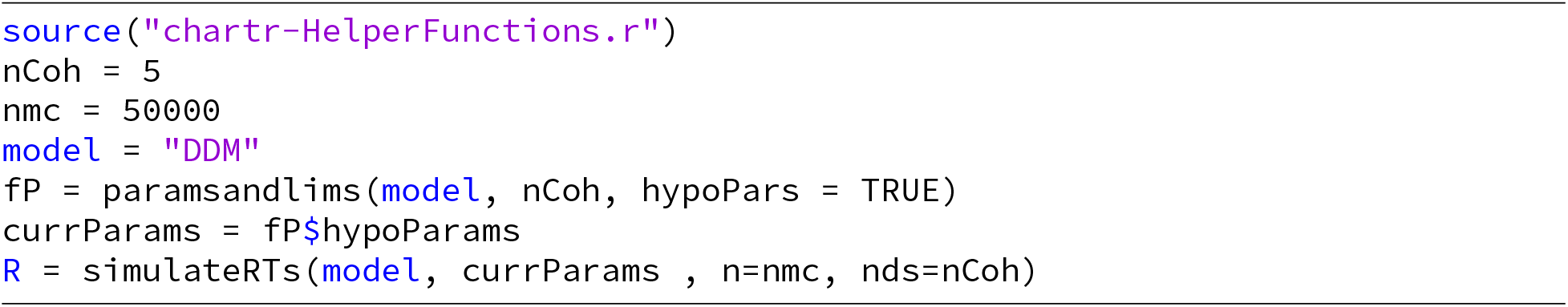

Fig. 5 shows the output of “chartr-Demo.r”, which simulates and visualizes choice and RT data from four models in *CHaRTr* : DDM, DDMS_v_S_z_S_t_, UGMS_v_, and dDDMS_v_. Fig. 5A shows predictions of the simple DDM (see Section 2.1.1), a symmetric, inverted-U shaped QP plot (Ratcliff and McKoon, 2008); the symmetry implies that correct and error RTs are identically distributed. As variability is introduced to the DDM’s starting state (*S*_*z*_) and/or drift-rate (*S*_*v*_; see Section 2.1.2), the QP plot loses its symmetry (Fig. 5B); relative to correct RTs, error RTs can be faster (due to *S*_*z*_) or slower (due to *S*_*v*_). Fig. 5B also introduced variability in non-decision time (*S*_*t*_), which increases the variance of the fastest responses.

**Figure 5:**
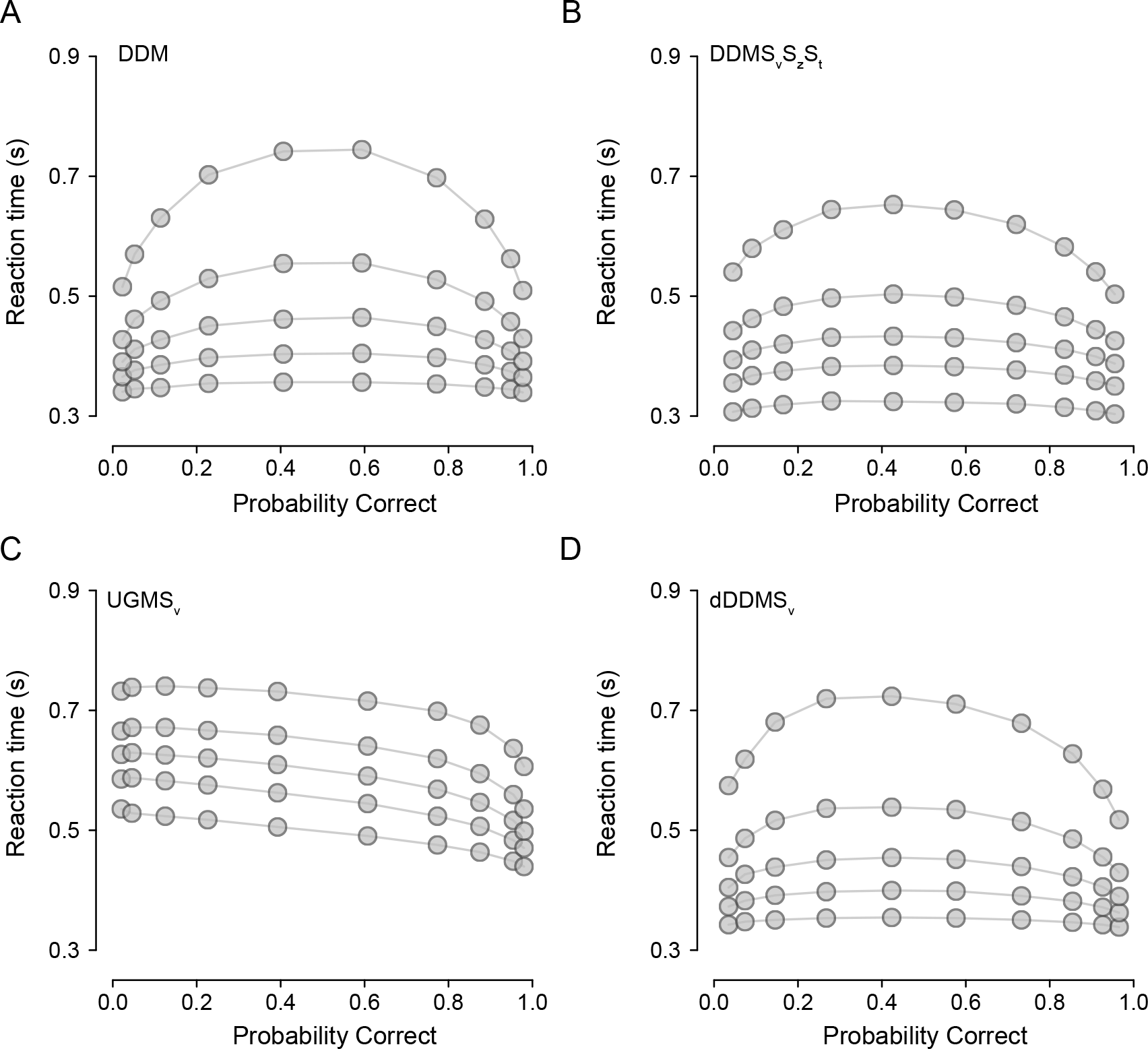
Quantile probability plots of data simulated from four models in *CHaRTr*. (A) DDM, (B) DDM with variable drift rates, starting state and non-decision time (DDMS_v_S_z_S_t_), (C) Urgency gating model with variable drift rates (UGMS_v_), and (D) DDM with an urgency signal defined as per Ditterich (2006a) (dDDMS_v_). Gray points denote data. Lines are drawn for visualization purposes.

Fig. 5C shows predictions of a standard variant of the UGM model (UGMS_v_) that assumes variable drift rate, zero intercept, a slope (*β*) of 1 and a time constant of 100 ms (see Section 2.1.5). The urgency gating mechanism in this model reduces the positive skew of the RT distributions, and leads to the prediction that error RTs are always slower than correct RTs (Fig. 5C; Hawkins et al., 2015b). Like the UGM, the dDDMS_v_ model, another model of urgency (see Section 2.1.4), also predicts reduced positive skew of the RT distributions. Unlike the standard UGM, however, it can also predict error RTs that are faster or slower than correct RTs (Fig. 5D).

It is clear from Fig. 5 that various features in data discriminate between various features of the decision-making models: the relative speed of correct and error RTs, and critically the shape of complete RT distributions. We now provide three illustrative case studies that take advantage of the differential predictions of the models, demonstrating the use of *CHaRTr* for model parameter estimation and selection amongst sets of competing models.

### 3.4. Case Studies

To illustrate the utility of the toolbox, we provide three case studies where we simulated data from decision-making models in *CHaRTr* (case studies 1 and 2) or use *CHaRTr* to model data collected from monkeys performing a decision-making task (case study 3). We use the case studies to demonstrate the typical model estimation and selection analyses. The case studies also provide a test of model and parameter recovery. That is, whether *CHaRTr* reliably selects the true data-generating model, and whether it reliably estimates the parameters of the true data-generating model.

#### 3.4.1 Case Study 1: Hypothetical Data Generated from a DDM with Variable Drift Rate and Non-Decision Time (DDMS_v_S_t_)

For our first case study we assumed the data came from hypothetical observers who made decisions in a manner consistent with a DDM with variable drift rate (*S*_*v*_) and variable start times (*S*_*t*_). In *CHaRTr*, this corresponds to simulating data from the model DDMS_v_S_t_, where an observer’s RTs exhibit variability due to both the decision-formation process and the non-decision components. We simulated 300 trials for each of 5 stimulus difficulties, for 5 hypothetical participants.

For each model and hypothetical participant, we repeated the parameter estimation procedure 5 times, independently. We heavily recommend this redundant-estimation approach as it greatly reduces the likelihood of terminating the optimization algorithm in local minima, which can arise in simulation-based models like those implemented in *CHaRTr*. Variability occurs due to randomness in simulating predictions of the model at each iteration of the optimization algorithm, and randomness in the optimization algorithm itself (for similar approach, see Hawkins et al., 2015a,b). We then select the best of the 5 independent parameter estimation procedures (or ‘runs’) for each model and participant (i.e., the ‘run’ with the highest value of the QMP statistic). If computational constraints are not an issue, then we encourage as many repetitions as possible of the parameter estimation procedure.

Fig. 6A shows the BICs for a set of models, obtained after using *CHaRTr* to fit the RT and choice data from one of the hypothetical observers. All the BICs are reported with reference to the DDM (i.e., as difference scores relative to the DDM). Thus, negative values of the BIC score suggest a more parsimonious account of the data than the DDM, and positive values suggest the opposite.

**Figure 6:**
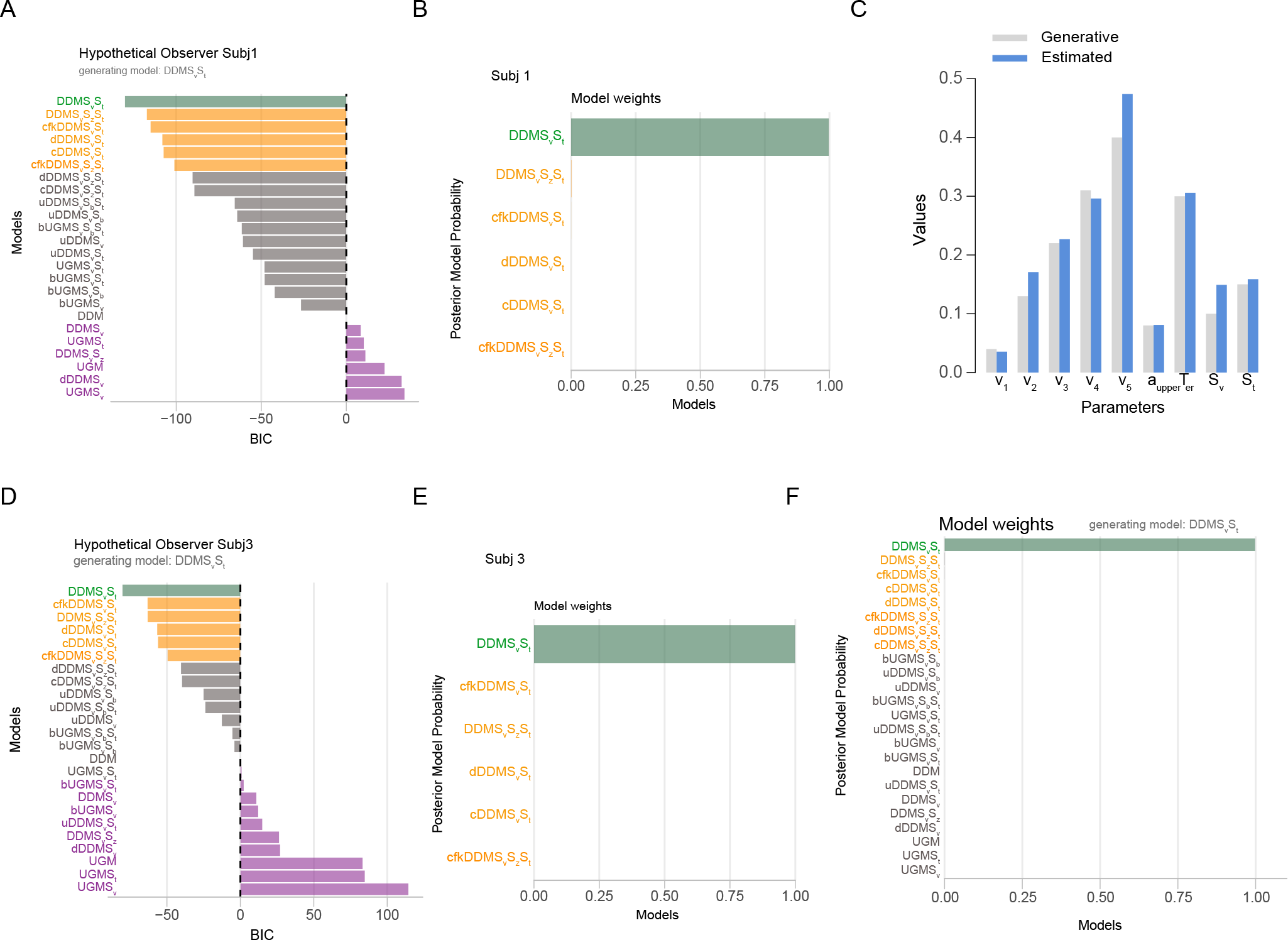
Model selection and parameter estimation outcomes from applying a range of cognitive models of decision-making to hypothetical data from two observers (case study 1). A-C shows outcomes from one hypothetical observer. D-E shows outcomes from a second hypothetical observer. Data were generated using the model DDMS_v_S_t_. A) BIC values as a function of model with the DDM model as the reference. B) Akaike weights for the top six models that provided the best account of the data. *CHaRTr* correctly identifies the true data-generating model (DDMS_v_S_t_) as the most likely candidate for describing the data. C) Data-generative and estimated parameter values for the DDMS_v_S_t_ model shown in A. Close alignment indicates *CHaRTr* recovered the true parameter values. D-E shows the BIC values and posterior model probabilities from another hypothetical observer. F) Shows the average posterior model probabilities across all five hypothetical observers, assuming them to be independent. Reassuringly, DDMS_v_S_t_ is identified as the most plausible model for the data.

Fig. 6B shows the BIC-based approximate posterior model probabilities (Eq. 21) for the top six models. DDMS_v_S_t_ provided the best account of the data; by ‘best account’, we mean the model that provided the most appropriate tradeoff between model fit and model complexity among the specific set of models under consideration, according to BIC. This suggests that *CHaRTr* can successfully identify the data-generating model – a necessary test for any parameter estimation and model selection analysis. We strongly recommend this form of *model recovery* analysis when developing and testing any proposed cognitive model; if a candidate model cannot be successfully identified in simulated data, where the true model is known, it will not be useful a model for real data.

The models *CHaRTr* ranked 2^nd^ to 6^th^ were sensibly related to the data-generating model. These models all assumed that observed RTs were influenced by factors other than sensory evidence (such as growing impatience), which might mimic the data-generating model’s RT variability that arose due to factors external to the decision-formation process (variable non-decision time). Although these results indicate that the DDMS_v_S_t_ model provided a better account of the data than cfkDDMS_v_S_t_, dDDMS_v_S_t_, cDDMS_v_S_t_, and DDMS_v_S_z_S_t_, they also serve as an important reminder that model selection should not be used to argue for the “best” model in an absolute sense. Rather, it is often most constructive to rank useful hypotheses/explanations about the data that can then guide further study (Burnham et al., 2011), which is the approach we have used here.

Fig. 6C shows the estimated parameter values for the best-fitting DDMS_v_S_t_ model. The parameter estimates were very similar to the data-generating values, with some minor over-or under-estimation of the drift rate parameters. This suggests that *CHaRTr* can reasonably recover the data-generating model *and* parameters. As above, we also strongly recommend this form of *parameter recovery* analysis when developing and testing any proposed cognitive model.

Fig. 6D and Fig. 6E show the model selection outcomes from another hypothetical observer. The best fitting model is again identified as DDMS_v_S_t_. A few other models also provided good accounts of the data. As was the case for observer 1, these models predict variability in RTs due to mechanisms outside the decision-formation process.

In the three other hypothetical observers that we simulated, the pattern of results returned by *CHaRTr* was consistent with the results shown for the two hypothetical observers in Fig. 6: DDMS_v_S_t_ was chosen as the best fitting model for all observers. If we assume the set of observers are independent (which they are in the case of our hypothetical example and usually in experiments), we can average over their posterior model probabilities to obtain a group-level estimate. As shown in Fig. 6F, DDMS_v_S_t_ is identified as the most plausible model for the data across the set of observers, indicating good model recovery.

Fig. 7 shows QP plots of the data from two hypothetical observers overlaid on the predictions from a range of models. The simple DDM predicted larger variance than was observed in data, and therefore provided a poor account of the data. When the DDM is augmented with S_v_ and S_t_, it provided a much improved account of the data, capturing most of the RT quantiles and the accuracy patterns. Three other models provided an almost-equivalent account of the data in terms of log-likelihoods (DDMS_v_S_z_S_t_, cDDMS_v_S_t_, dDDMS_v_S_t_), but they did so with the use of more model parameters than DDMS_v_S_t_. This led to a larger complexity penalty for those models and thus larger BICs in comparison to the DDMS_v_S_t_ model, as shown in the model selection analysis in Fig. 6.

**Figure 7:**
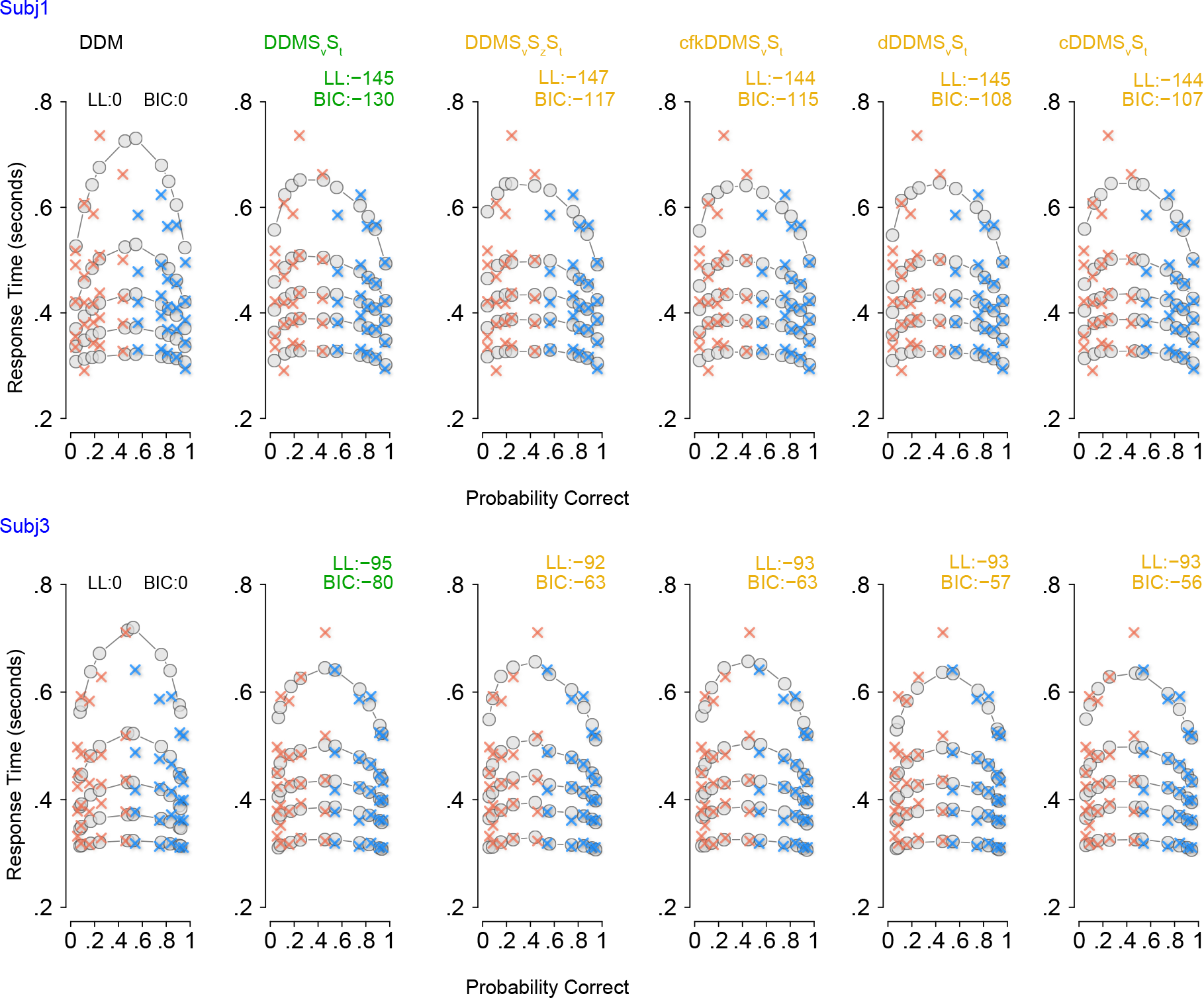
Quantile probability (QP) plots showing correct RTs (blue) and error RTs (orange) for two hypothetical observers (case study 1), along with the model predictions (gray dots). Predictions from the four best-fitting models are shown along with the simplest model the DDM. The four best fitting models are DDMS_v_S_t_, cfkDDMS_v_S_t_, dDDMS_v_S_t_, cDDMS_v_S_t_ are shown. Numbers at the top of each plot show the BIC for the model under consideration, assuming the DDM as the base (reference) model. The model DDMS_v_S_t_provides the best account of the data.

### Case Study 2: Hypothetical Data Generated from a UGM with Variable Intercept (bUGMS_v_)

In a second case study we simulated data from hypothetical observers whose decision-formation process was controlled by an urgency gating model (UGM) with a variable drift rate and an intercept (Cisek et al., 2009; Thura et al., 2012), termed bUGMS_v_ in *CHaRTr*. We again assumed five hypothetical subjects, five stimulus difficulties and simulated 500 trials for each of them. We then fit the data with the redundant-estimation approach as in case study 1 and evaluated the results of the model selection analysis, all using routines contained in *CHaRTr*.

Fig. 8A shows the BICs for the set of models considered for one hypothetical observer’s data, again referenced to the DDM (i.e., as difference scores relative to the DDM). Negative values of the BIC score suggest a more parsimonious account of the data than the DDM, and positive values suggest the opposite. Fig. 8B shows the BIC weights (Eq. 21) for the top six models. bUGMS_v_ provided the best account of the data for this hypothetical observer. The models *CHaRTr* ranked 2^nd^ to 6^th^ were also sensibly related to the data-generating model; they all assumed the decision-formation process was influenced by factors other than sensory evidence, such as growing impatience. The second case study reaffirms our conclusion from the first case study that model selection may not be put to best use when arguing for a single “best” model in an absolute sense. This is especially true when the data-generating model is not decisively recovered from data.

**Figure 8:**
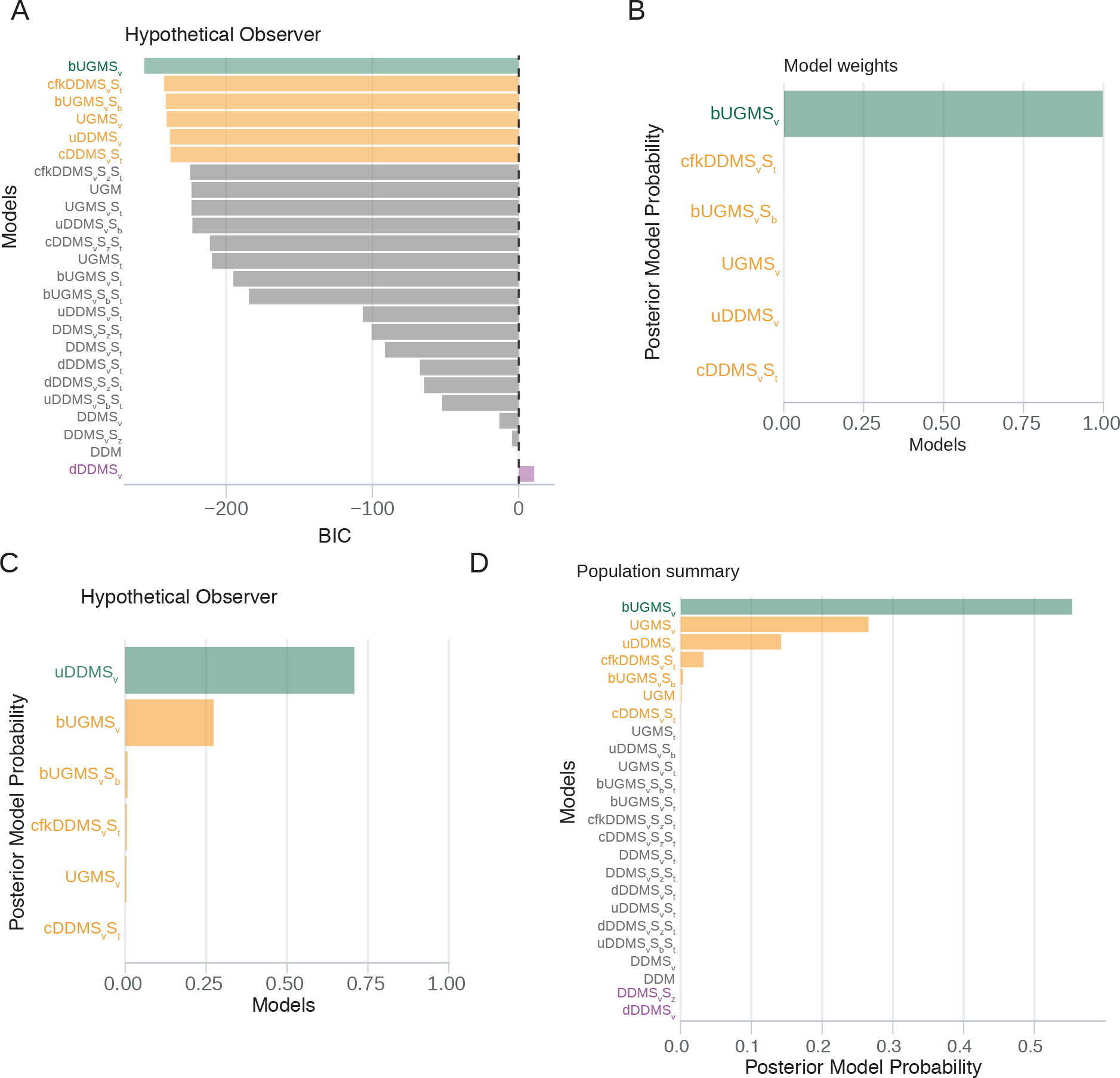
Model selection and parameter estimation outcomes from applying a range of cognitive models of decision-making to data from a hypothetical observer. Decision-making in this hypothetical observer is controlled by the model bUGMS_v_. A) BIC values as a function of model with the DDM model as the reference for one hypothetical observer, Subj 3. B) Posterior model probabilities for the top six models that provided the best account of Subj 3’s behavior. C) Results for another hypothetical subject. D) Results for the population of hypothetical subjects. The most probable model for this set of hypothetical observers is the generative model, bUGM S_v_. However, we note that other models such as UGMS_v_, and uDDMS_v_ provide quite good descriptions of the behavior. This result is in keeping with the general notion that model selection ought to be used as a guide to the most likely models and not necessarily to argue for a “best” model.

Model selection sometimes fails to recover the data-generating model. Fig. 8C shows the top six models identified by *CHaRTr* as providing the best fit to another of the hypothetical observers’ data; uDDMS_v_ provided a better fit than the generative model bUGMS_v_. This result highlights two important points. First, some models under some circumstances can mimic each other (i.e., generate similar predictions), which makes their identification in data difficult. Second, some models may not be mimicked, but they may require very many data points to reliably recover. We note that these points are not specific to *CHaRTr* – they are properties of quantitative model selection in general and are an important reminder of the necessary careful steps needed when aiming to select between models (Chandrasekaran et al., 2018).

Fig. 8D shows the posterior model probabilities for the different models averaged over all five observers considered. Reassuringly, the most plausible model across the set of observers is the generative model bUGMS_v_. The next five best models are all conceptually related to the data generating model. For instance, the next best model was UGMS_v_ which is an urgency gating model with no intercept. The third best model was uDDMS_v_ which is a DDM model with urgency but no gating. Together these results again serve as a reminder of the utility of *CHaRTr* in the analysis of decision-making models, including the ability to quantitatively assess a large set of conceptually similar and dissimilar models.

#### 2.4.3 Case Study 3: Behavioral Data From Monkeys Reported in Roitman and Shadlen (2002)

To demonstrate the utility of *CHaRTr* in understanding experimental data, we use *CHaRTr* to model the freely available RT and choice data from two monkeys performing a random-dot motion RT decision-making task (Roitman and Shadlen, 2002). In this classic variant of the random-dot motion task, the monkeys were trained to report the direction of coherent motion with eye movements. The percentage of coherently moving dots was randomized from trial to trial across six levels (0%, 3.2%, 6.4%, 12.8%, 25.6% and 51.2%). Monkey b completed 2614 trials and Monkey n completed 3534 trials.

We demonstrate that *CHaRTr* replicates key findings from past analyses of these behavioral data. Roitman and Shadlen (2002)’s behavioral (and neural) data were originally interpreted as a neural correlate of the DDM. Later studies suggested a stronger role for impatience/urgency in these data (Ditterich, 2006b; Hawkins et al., 2015b). This is the first result we wish to demonstrate again using *CHaRTr*. Second, Hawkins et al. (2015a) showed that the urgency gating model provides a better description of the data than the DDM. We note that recent work suggests the evidence for impatience/urgency in Roitman and Shadlen (2002)’s data might be the result of the particular training regime the monkeys were exposed to (Evans and Hawkins, 2019).

Fig. 9A-B shows the results from *CHaRTr*. For both monkeys, the four best-performing models all included a DDM with either urgency or collapsing bounds, and the worst performing models were largely DDM models without any forms of urgency. As mentioned above, for any functional form of a collapsing boundary there is a form of additive urgency signal that can generate identical predictions. So finding that collapsing bound models describe the data better is consistent with prior observations that urgency is an important factor. Together, the results are broadly consistent with those of Ditterich (2006a) and Hawkins et al. (2015b) who reported that models with forms of impatience are systematically better than models without it, for Roitman and Shadlen (2002)’s data. Fig. 9C-D shows that when the comparison is restricted to UGM and DDM models, variants of the UGM better explain the behavior of the monkeys than variants of the DDM, which is consistent with the findings of Hawkins et al. (2015a).

**Figure 9:**
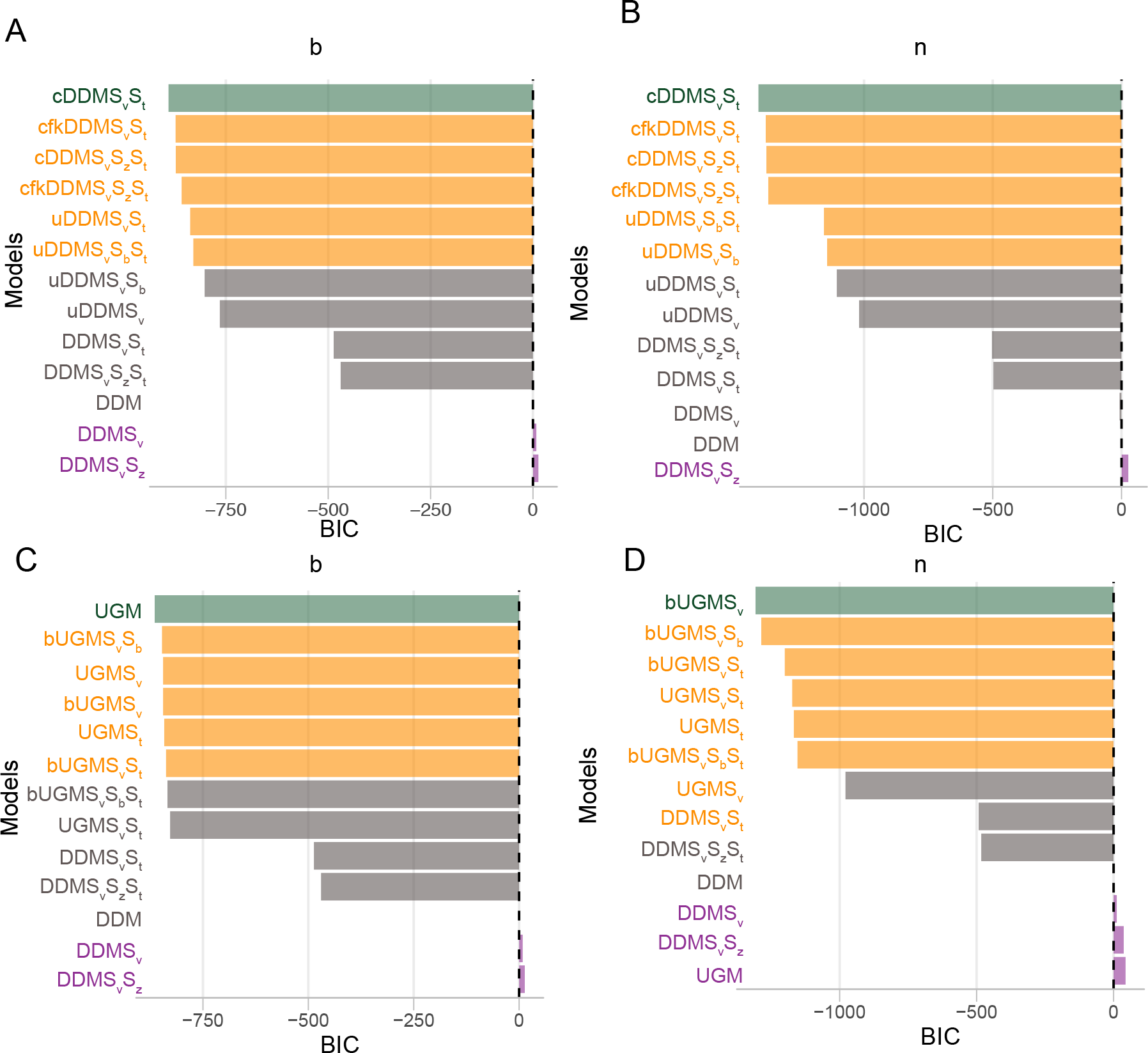
Model selection outcomes from applying a range of cognitive models of decision-making to data from two monkeys (Roitman and Shadlen, 2002). A-B shows outcomes from monkey b and n to compare models with various forms of urgency vs. simple diffusion decision models without impatience. For both monkeys, *CHaRTr* suggests models with urgency are better candidates for describing the data than DDMs without urgency. C-D shows outcomes from the monkeys b and n when comparing UGM vs. DDM models. For both monkeys, UGM based models substantially outperform the DDM based models

We can take things one step further and use *CHaRTr* to derive more insights into the behavior of the monkeys in this decision-making task, by examining whether urgency or the time constant of integration is a more important factor in explaining their behavior. Fig. 10 shows quantile probability plots for five models: DDMS_v_S_z_S_t_, a model from the DDM class without urgency but elaborated with variability in various parameters (S_v_, S_z_, S_t_), two models with Urgency and variability in some parameters (uDDMS_v_S_t_, uDDMS_v_S_b_), and two UGM models with variability in parameters (bUGMS_v_S_b_, bUGMS_v_S_t_). As shown in Fig. 9, addition of urgency dramatically improved the ability of these models to account for the decision-making behavior of the two monkeys. We next used *CHaRTr* for a preliminary analysis of whether the gating component of the urgency gating model improves model predictions over and above urgency alone. In both monkeys, we found that the data are slightly more consistent with models such as bUGMS_v_S_b_ and bUGMS_v_S_t_, models that involve urgency and gating with a 100 ms time constant of integration. These observations provide hypotheses for further analyses of the neural data and further targeted model selection.

**Figure 10:**
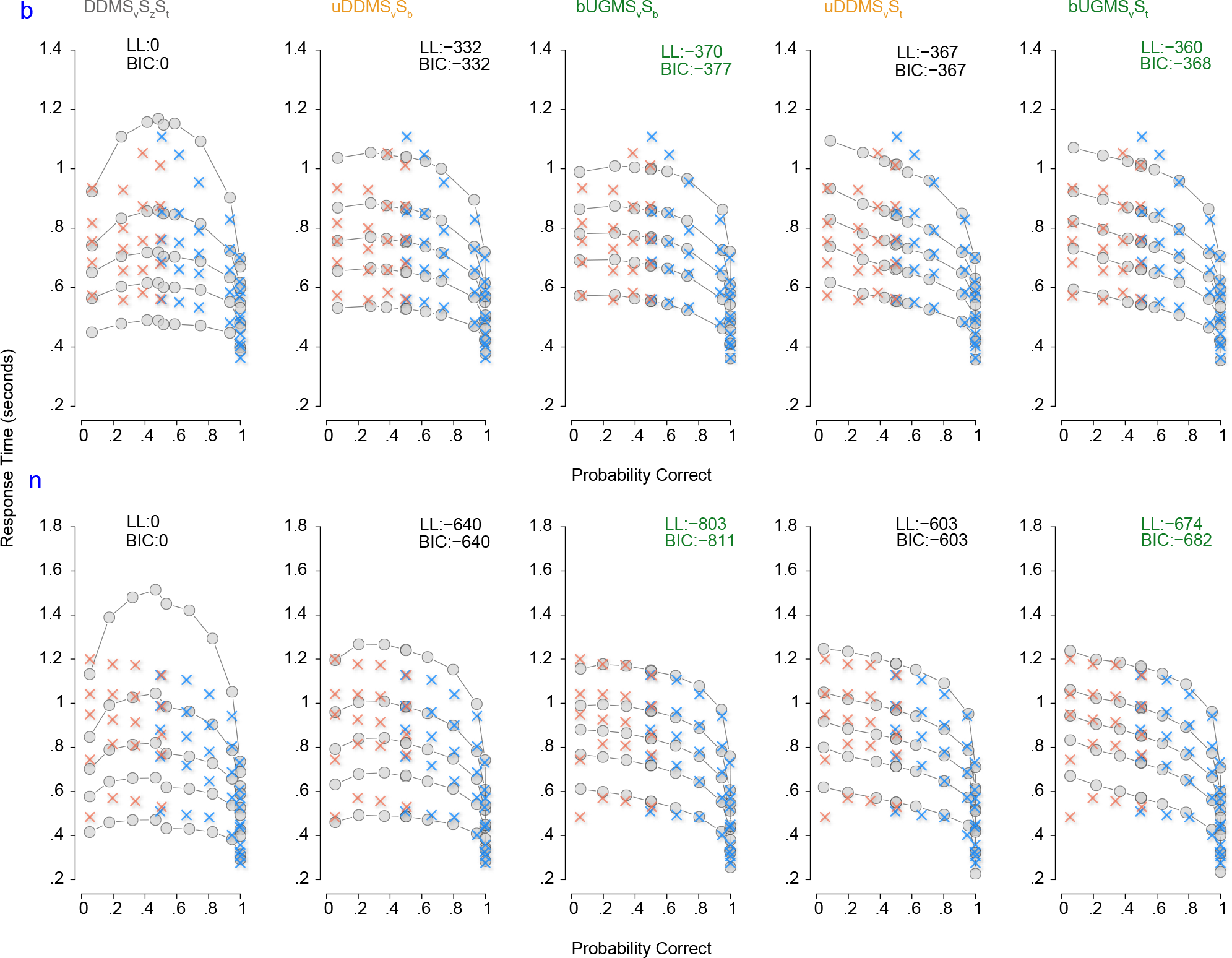
Quantile probability (QP) plots showing data in blue (corrects) and yellow crosses (errors) for the two monkeys from Roitman and Shadlen (2002), along with the model predictions (gray dots). Predictions from DDMS_v_S_z_S_t_ are shown along with four other models uDDMS_v_S_b_, bUGMS_v_S_b_, uDDMS_v_S_t_, bUGMS_v_S_b_. Numbers at the top of each plot show the BIC for the model under consideration, assuming DDMS_v_S_z_S_t_ as the base (reference) model. For both monkeys the model bUGMS_v_S_b_ is the best model for describing the data out of these candidate set of models

Together, the results in Fig. 9 and Fig. 10 highlight the ease with which *CHaRTr* can be used to make insightful statements about behavior in decision-making tasks and ultimately may be a stepping stone for deeper insights into mechanism (Krakauer et al., 2017).

## 4. Discussion

Advances in our understanding of decision making have come from three fronts: 1) through novel experimental manipulations of sensory stimuli (Brody and Hanks, 2016; Cisek et al., 2009; Ratcliff, 2002; Ratcliff and Rouder, 2000; Smith and Ratcliff, 2009; Thura and Cisek, 2014) and/or task manipulations (Hanks et al., 2014), 2) recording neural data in a variety of decision-related structures in multiple model systems (Chandrasekaran et al., 2017; Coallier et al., 2015; Ding and Gold, 2012a; Hanks et al., 2015; Schall, 2001; Shadlen and Newsome, 2001; Thura et al., 2014) and 3) developing and testing quantitative cognitive models of choices, RTs, and other behavioral readouts from animal and human observers performing these decision-making tasks (Ratcliff and Smith, 2015). Quantitative modeling is a lynchpin in generating novel insights into cognitive processes such as decision-making. However, it has posed significant technical and computational challenges to the researcher. Widespread and rapid uptake of quantitative modeling requires software toolboxes that can easily implement the many sophisticated models of decision-making proposed in the literature (Diederich, 1997b; Ratcliff et al., 2016; Thura et al., 2012).

We contend that the ideal toolbox for developing and implementing cognitive models of decision-making and evaluating them against choice and RT data should be simple, offer a plurality of cognitive models, provide model estimation and model selection procedures, provide simple simulation and visualization tools, and be easily extensible when new hypotheses are developed. Such a view is broadly consistent with recent research that lays out the best practices for computational modeling of behavior (Heathcote et al., 2015; Wilson and Collins, 2019). Ready adoption is also facilitated when the toolbox is implemented in an open-source, free programming language obviating the need for expensive licenses. The added benefit of an open source toolbox is that researchers can look “under the hood”, which has at least three benefits: 1) allow a deeper level of understanding of the models, 2) readily permit extension of the toolbox, and 3) catch errors in implementation. At the time of development of this toolbox and submission of this study, no existing toolbox has satisfied all of these criteria.

*CHaRTr* was guided by these pragmatic principles, and is our attempt to provide a practical toolbox that encompasses a range of cognitive models of decision-making. Some of the models are grounded in classic random walk and diffusion models (Ratcliff, 1978; Stone, 1960). Others incorporate modern hypotheses that decision-making behavior might involve signals such as urgency (Ditterich, 2006b), collapsing boundaries (Drugowitsch et al., 2012), and variable non-decision times (Ratcliff and Tuerlinckx, 2002). Since all of the source code is freely available, the toolbox thus provides a framework where models that are proposed into the future can also be implemented and contrasted against existing models. We provide a suite of functions for estimating the parameters of decision-making models, methods to compare log-likelihoods, and calculating penalized information criteria from these different models. Finally, the toolbox is developed in the R Statistical Environment, an open source language that is maintained by an active community of scientists and statisticians (R Core Team, 2016).

We anticipate that *CHaRTr* will provide a pathway to standardizing quantitative comparisons between models and across studies, and ultimately serve as one of the reference implementations for researchers interested in developing and experimentally testing candidate models of decision-making processes. *CHaRTr* also codifies the various parameters of decision-making models, which reflects the hypothesized latent constructs and how they interact, and provides easy access to more than 20 models of behavioral performance in decision-making tasks including variants of the diffusion decision model, the urgency gating model, diffusion models with urgency signals, and diffusion models with collapsing boundaries. *CHaRTr* also offers pedagogical value because it allows the user to effortlessly simulate the many different models of decision-making and generate RT and choice data from hypothetical observers. *CHaRTr* will also allow quantitative evaluation of the predictions of various decision-making models and help move away from qualitative intuition-based predictions from these models. Finally, *CHaRTr* is also sufficiently flexible that users can implement novel models with their own specific assumptions.

*CHaRTr* provides researchers with the resources to apply and test more than 20 different, albeit overlapping, variants of decision-making models. We have argued throughout that model selection techniques ought to be used as a tool for selecting families of models to guide the next generation of experiments and further analyses, which is in the spirit of Burnham et al. (2011); we do not believe model selection should be used to justify categorical answers ("the best model"). In this sense, model selection is one tool in the whole gamut of tools that are needed to understand decision-making (Chandrasekaran et al., 2018)

The most promising approaches for advancing our understanding of decision-making will combine the rigorous model selection techniques we advocate here with novel experimental manipulations of stimulus statistics (Brody and Hanks, 2016; Cisek et al., 2009; Evans et al., 2017; Thura et al., 2014), task contingencies (Hanks et al., 2014; Heitz and Schall, 2012; Murphy et al., 2016; Thura and Cisek, 2016), and a range of other factors. We believe that validating and advancing models of decision-making will be facilitated by data that is freely available for the kinds of model estimation and model selection analyses we have performed here. Here, we took advantage of the freely available dataset from Roitman and Shadlen (2002). We anticipate the application of *CHaRTr* to many more decision-making datasets will help to form a coherent picture of how various latent cognitive processes affect the behavior of animal and human decision-making. This deeper understanding of decision-making behavior (Krakauer et al., 2017) will in turn facilitate a deeper understanding of decision-related neural responses (Chandrasekaran et al., 2017; Churchland et al., 2008; Cisek et al., 2009; Murphy et al., 2016; O’Connell et al., 2018; Purcell and Kiani, 2016; Thura et al., 2012).

Rigorous model selection techniques are even more relevant if we wish to make further inroads into understanding the neural correlates of decision-making. In particular, discriminating between multiple candidate models of decision-making is critical for neurophysiological studies of decision-making that attempt to relate neural responses in decision-related structures to the features of sequential sampling models (Ditterich, 2006a,b; Gold and Shadlen, 2007; Hanes and Schall, 1996; Heitz and Schall, 2012; Shadlen and Newsome, 2001; Thura et al., 2012). For example, one of the most well-established tenets of the neural basis of decision-making is the gradual ramp-like increase in the firing rates of individual neurons in decision-related structures such as the lateral intraparietal area (Roitman and Shadlen, 2002; Shadlen and Newsome, 2001), frontal eye fields (Ding and Gold, 2012a; Hanes and Schall, 1996), superior colliculus (Ratcliff et al., 2003, 2007), prefrontal cortex (Kim and Shadlen, 1999) and dorsal premotor cortex (Chandrasekaran et al., 2017; Coallier et al., 2015; Thura et al., 2014). However, questions still remain; for example, is the ramp in a neuron’s response a signature of the evidence integration process posited by a DDM or is it more consistent with the presence of, say, an increasing urgency signal. It can be challenging to neurally discriminate between frameworks without a clear hypothesis about 1) a detailed and ideally quantitative understanding of the behavior (Krakauer et al., 2017; O’Connell et al., 2018), and 2) the mapping from the underlying neural mechanisms to the observed behavior (Schall, 2004). We believe *CHaRTr* and other toolboxes of its ilk will play a critical role in further advancing our understanding of the neural correlates of decision-making.

### 4.1. Future directions

*CHaRTr* provides a powerful framework for estimating and discriminating between candidate decision-making models. Nevertheless, there is considerable scope for extending its capabilities. Here, we outline a few future directions we believe would make *CHaRTr*, and other toolboxes that come in its wake, even more useful for decision-making researchers.

First, *CHaRTr* provides options to estimate sequential sampling models that assume *relative* evidence is accumulated over time. A related and compelling line of research assumes a race model architecture where a choice between *n* options is represented as a race between *n* evidence accumulators. The *n* ≥ 2 accumulators collect evidence in favor of their respective response options as a dynamic race toward their respective thresholds. The first accumulator to reach the threshold triggers a decision for the corresponding response option. There are a range of race models that differ in details, including accumulators that are independent (e.g., Brown and Heathcote, 2008; Reddi and Carpenter, 2000) or dependent (e.g., Usher and McClelland, 2001). Naturally, these models can be elaborated with many features of the *relative* evidence accumulation models implemented in *CHaRTr*, including variable non-decision times and urgency (though see Bogacz et al., 2006; Zhang et al., 2014, for demonstration of the equivalence between relative and absolute evidence accumulation models under certain circumstances). Incorporation of race models in *CHaRTr* will be a useful extension into the future.

Second, the current instantiation of *CHaRTr* assumes that observers are independent. Recent efforts have proposed the use of hierarchical Bayesian methods for the DDM and other decision-making models (Ahn et al., 2017; Heathcote et al., 2018; Wiecki et al., 2013). Bayesian methods provide two advantages over the current framework provided in *CHaRTr*. Bayesian methods of parameter estimation incorporate prior knowledge into the plausible distribution of parameter values, and provide full posterior distributions for all model parameters. *CHaRTr* currently provides only the most likely value for a parameter without any measure of its uncertainty, whereas the full posterior distribution provides uncertainty in the estimate for each parameter, thus reducing the likelihood of drawing over-confident conclusions. Bayesian methods are also advantageous when used in contexts where there are only modest numbers of trials per observer. Hierarchical Bayesian models in particular can enhance statistical power by providing opportunities for simultaneous estimation of the parameters of individual observers as well as the population-level distributions from which they are drawn.

Despite these benefits, we emphasize that it is far from straightforward to extend the models implemented in *CHaRTr* to Bayesian parameter estimation methods. The goal of *CHaRTr* is simple and rapid implementation and testing of new models, which takes place via simulation-based techniques. Bayesian methods require model likelihood functions, which can be challenging to derive and may not even exist for some of the models implemented in *CHaRTr*, and as such the extension to Bayesian methods is not trivial. In future work, we aim to extend the parameter estimation routines in *CHaRTr* to make use of approximate Bayesian techniques.

Third, the framework in *CHaRTr* is currently only amenable for analyzing behavior from decision-making tasks where the sensory stimulus provides constant evidence over time, albeit with noise, and varies along a single dimension. However, previous research suggests that a powerful way to dissociate between different models of decision-making is to use time-varying stimuli (Brody and Hanks, 2016; Brunton et al., 2013; Cisek et al., 2009; Ratcliff, 2002; Ratcliff and Rouder, 2000; Smith and Ratcliff, 2009; Thura et al., 2014; Usher and McClelland, 2001). In a related vein, there has been increased interest in combining frameworks that posit sensory stimuli are optimally combined and could drive multisensory decision-making models (Chandrasekaran et al., 2017; Drugowitsch et al., 2014). Future versions of *CHaRTr* will provide opportunities for implementing and testing models in contexts where the sensory stimuli have temporal structure (Evans et al., 2017), or involve multi-sensory integration (Chandrasekaran et al., 2017).

Finally, *CHaRTr* currently allows the quality of the evidence signal (drift rate) to vary with an experimental factor (stimulus difficulty). In future versions of *CHaRTr*, we will provide capabilities for different model parameters to vary with different experimental factors. There are a range of other experimental manipulations whose effect will likely appear in model parameters other than the drift rate; for example, emphasizing the speed or accuracy of decisions is most likely to affect the decision boundary, or the speed with which a boundary collapses. Future versions of *CHaRTr* will allow researchers to test and discriminate between these hypotheses.

## Acknowledgments

CC was supported by a NIH/NINDS R00 award 4R00NS092972-03. GH was supported by an Australian Research Council (ARC) Discovery Early Career Researcher Award (DECRA, award DE170100177) and an ARC Discovery Project (award DP180103613). Some of the model fits for simulated and real data were performed on the Shared Computing Cluster funded by an ONR DURIP N00014-17-1-2304 grant to Boston University. Some of the work was done under the auspices of Prof. Krishna Shenoy at Stanford University. We thank Prof. Shenoy for helpful discussions and advice, and Jessica Verhein and Megan Wang for insightful discussions.

